# *CACNA1A* haploinsufficiency leads to reduced synaptic function and increased intrinsic excitability

**DOI:** 10.1101/2024.03.18.585506

**Authors:** Marina P. Hommersom, Nina Doorn, Sofía Puvogel, Elly I. Lewerissa, Annika Mordelt, Ummi Ciptasari, Franziska Kampshoff, Lieke Dillen, Ellen van Beusekom, Astrid Oudakker, Naoki Kogo, Monica Frega, Dirk Schubert, Bart P.C. van de Warrenburg, Nael Nadif Kasri, Hans van Bokhoven

**Affiliations:** Department of Human Genetics, Radboud University Medical Center, Donders Institute for Brain, Cognition, and Behaviour, 6500 HB Nijmegen, The Netherlands; Department of Clinical Neurophysiology, University of Twente, 7522 NB Enschede, The Netherlands; Department of Cognitive Neurosciences, Radboud University Medical Center, Donders Institute for Brain, Cognition, and Behaviour, 6500 HB Nijmegen, The Netherlands; Department of Neurology, Radboud University Medical Center, Donders Institute for Brain, Cognition, and Behaviour, 6500 HB Nijmegen, The Netherlands

**Keywords:** *CACNA1A*, induced pluripotent stem cells, neuronal networks, micro-electrode array, human disease modeling

## Abstract

Haploinsufficiency of the *CACNA1A* gene, encoding the pore-forming α1 subunit of P/Q-type voltage-gated calcium channels, is associated with a clinically variable phenotype ranging from cerebellar ataxia, to neurodevelopmental syndromes with epilepsy and intellectual disability.

To understand the pathological mechanisms of *CACNA1A* loss-of-function variants, we characterized a human neuronal model for *CACNA1A* haploinsufficiency, by differentiating isogenic induced pluripotent stem cell lines into glutamatergic neurons, and investigated the effect of *CACNA1A* haploinsufficiency on mature neuronal networks through a combination of electrophysiology, gene expression analysis, and *in silico* modeling.

We observed an altered network synchronization in *CACNA1A*^+/−^ networks alongside synaptic deficits, notably marked by an augmented contribution of GluA2-lacking AMPA receptors. Intriguingly, these synaptic perturbations coexisted with increased non-synaptically driven activity, as characterized by inhibition of NMDA and AMPA receptors on micro-electrode arrays. Single-cell electrophysiology and gene expression analysis corroborated this increased intrinsic excitability through reduced potassium channel function and expression. Moreover, we observed partial mitigation of the *CACNA1A*^+/−^ network phenotype by 4-aminopyridine, a therapeutic intervention for episodic ataxia type 2.

In summary, our study pioneers the characterization of a human induced pluripotent stem cell-derived neuronal model for *CACNA1A* haploinsufficiency, and has unveiled novel mechanistic insights. Beyond showcasing synaptic deficits, this neuronal model exhibited increased intrinsic excitability mediated by diminished potassium channel function, underscoring its potential as a therapeutic discovery platform with predictive validity.

## Introduction

Synaptic neurotransmission is dependent on proper functioning of the precisely orchestrated neurotransmitter release machinery. Voltage-gated calcium channels, in particular the P/Q-type channels, Ca_v_2.1, are an essential part of this machinery.^1–4^ The pore-forming subunit of Ca_v_2.1 is encoded by the *CACNA1A* gene. Gain-of-function variants in this gene are classically related to familial hemiplegic migraine type 1, loss-of-function variants to episodic ataxia type 2, and CAG-repeat expansions to spinocerebellar ataxia type 6. Next to these classic entities, additional phenotypes have been associated with *CACNA1A* variants, leading to a broad *CACNA1A* disease spectrum.^5,6^ We and others have shown that *CACNA1A* heterozygous loss-of-function variants not only lead to the classical entity of ataxia, but also a range of other phenotypes including epilepsy and intellectual disability.^5,7^ Thus, *CACNA1A* haploinsufficiency results in a clinically variable phenotype that can also include neurodevelopmental features.

Mouse models with *Cacna1a* loss-of-function variants recapitulate the range of human clinical phenotypes, exhibiting a combination of ataxia, paroxysmal dyskinesia, absence epilepsy, and cognitive deficits.^8–13^ The motor incoordination of these models, as well as the high expression of *CACNA1A* in the cerebellum, has led to a thorough investigation of Ca_v_2.1 in cerebellar granule cells and Purkinje cells. At the cellular level, loss-of-function variants lead to a reduced current density of Ca_v_2.1, loss of Purkinje cell pace-making activity, and Purkinje cell axonal swelling.^10–15^ However, most of these models only exhibit these cellular and behavioral phenotypes when variants are homozygous or compound heterozygous.^10–13^ Furthermore, whereas *CACNA1A* is widely expressed throughout the brain, only a few studies have touched upon the effects of loss-of-function of Ca_v_2.1 in the cerebral cortex.^16,17^

To elucidate the pathology of *CACNA1A* haploinsufficiency in a human context, we have generated isogenic iPSC lines with a monoallelic frameshift variant in exon 8 of *CACNA1A* via CRISPR/Cas9.^18^ These iPSCs were subsequently differentiated into cortical glutamatergic neurons.^19^ We characterized the effect of *CACNA1A* haploinsufficiency on mature neuronal networks. Through a combination of electrophysiology, gene expression analysis, and *in silico* modeling, we have uncovered cellular and synaptic mechanisms underlying dysfunction in *CACNA1A-*haploinsufficient networks. Specifically, we found a reduction in synaptic number and strength, but, counterintuitively, an increase of intrinsic excitability through reduced function and expression of potassium channels. Our findings thus establish an iPSC-derived neuronal model for *CACNA1A* haploinsufficiency that reveals novel mechanistic insights beyond Ca_v_2.1 function.

## Materials and methods

### Human iPSC lines and CRISPR/Cas9 editing of *CACNA1A*

The control line used in this study (UCSFi001-A, obtained from the Coriell Institute (GM25256, RRID: CVCL_Y803)) was reprogrammed from skin fibroblasts of a 30-year old healthy male. This line was used for CRISPR/Cas9 editing to generate isogenic lines with *CACNA1A* variants as previously described.^18^ In short, after targeting Cas9 towards exon 8 using an sgRNA (5’-ACGTGAGCTCAATGGGTACA-3’), the double strand break was repaired by non-homologous end joining, yielding two clones with a heterozygous one base pair insertion (NM_001127221.1: c.1174_1175insT) (Fig. 1A, Supplementary Fig. 1A). The generated hiPSC lines were validated for pluripotency markers and trilineage differentiation potential (STEMdiff^TM^ Trilineage Differentiation Kit, STEMCELL Technologies) by immunocytochemistry (Supplementary Materials and methods, Supplementary Fig. 1C-D). Off-target analysis was performed by sequencing the top 3 off-target sites of the sgRNA predicted by Benchling (Biology Software) and CRISPOR^20^ (Supplementary Fig. 1E). Both *CACNA1A*^+/−^ clones showed typical iPSC morphology, positive expression of pluripotency markers (OCT4, NANOG and SSEA4), could differentiate to all three lineages, and did not show any off-target alterations (see Hommersom *et al*. for *CACNA1A*^+/−^ (1)^18^ and Supplementary Fig. 1B-E for *CACNA1A*^+/−^ (2)). Copy number variation analysis by whole exome sequencing did not reveal any acquired variants in comparison to the published profile of this control line.^21^

**Figure 1.**
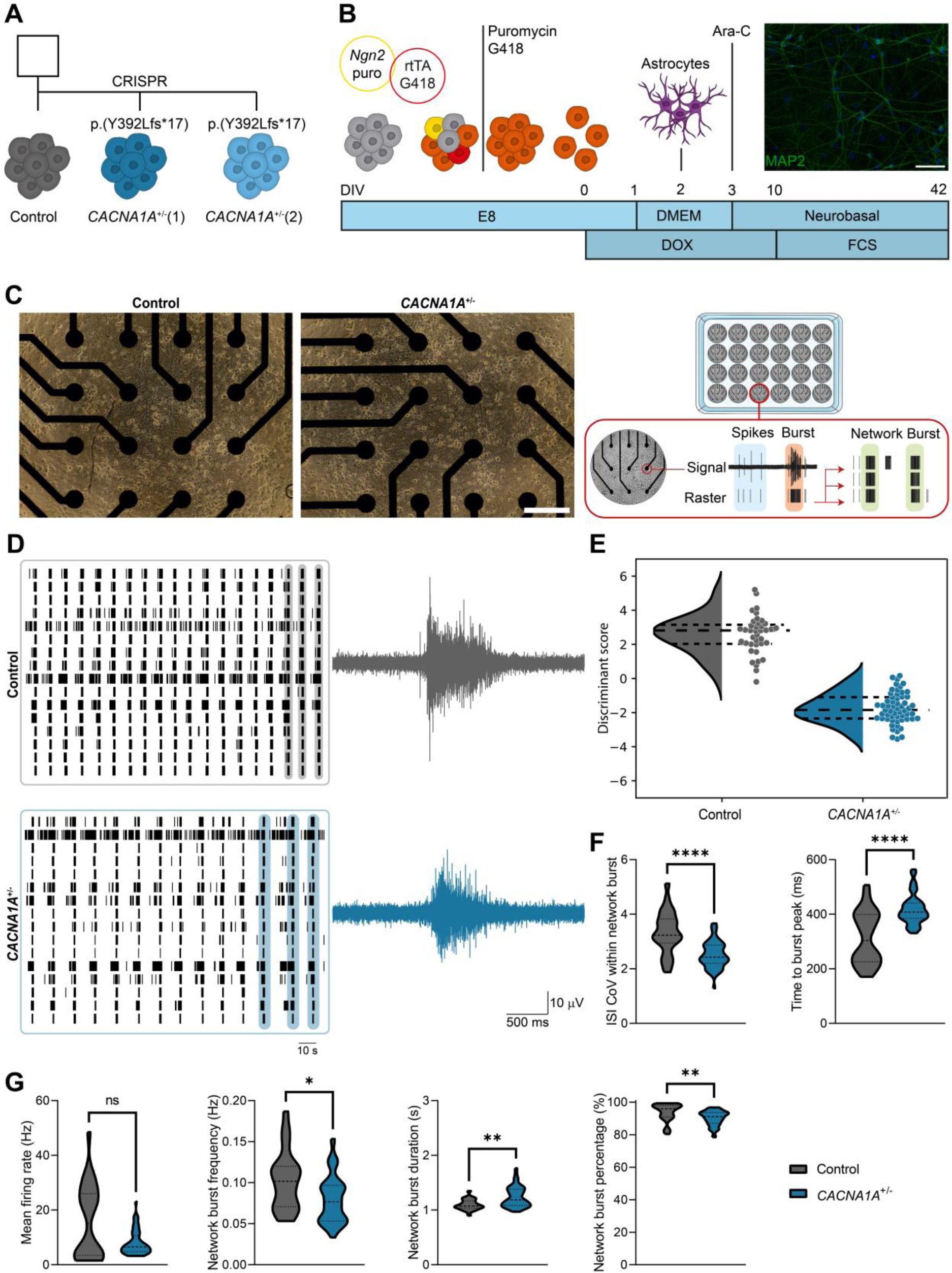
*CACNA1A*^+/−^ glutamatergic neurons display an altered network synchronization. (**A**) Schematic overview of the cell lines used in this study. (**B**) Schematic representation of the differentiation protocol from iPSCs to glutamatergic neurons, with a representative image of MAP2-positive control neurons at DIV42 (scale bar 100 μm). (**C**) Representative images of control and *CACNA1A*^+/−^ cells on MEA (left panel; scale bar 300 μm), with a schematic overview of electrophysiological activity from neurons cultured on MEA (right panel). (**D**) Representative 3-min rasterplots of spontaneous activity from control and *CACNA1A*^+/−^ networks at DIV42, with a 3-sec zoom-in of the activity of one electrode (bottom panel). (**E**) Discriminant scores of control and *CACNA1A*^+/−^ neurons based on discriminant analysis of eighteen MEA parameters as indicated in the structure matrix of Table 1. (**F**) Quantification of network parameters including inter spike interval (ISI) coefficient of variation (CoV) within network burst, and time to burst peak. (**G**) Quantification of network parameters including mean firing rate, network burst frequency, network burst duration, and network burst percentage (percentage of spikes within a network burst). *n = 24/4* for Control; *n = 57/4* for *CACNA1A*^+/−^ ((1): *29/4* and (2): *28/4*). Dashed line represents the median, dotted line represents the quartiles. *\*p < 0.05, **p < 0.01, ****p < 0.0001*, Mann–Whitney test.

**Table 1.**
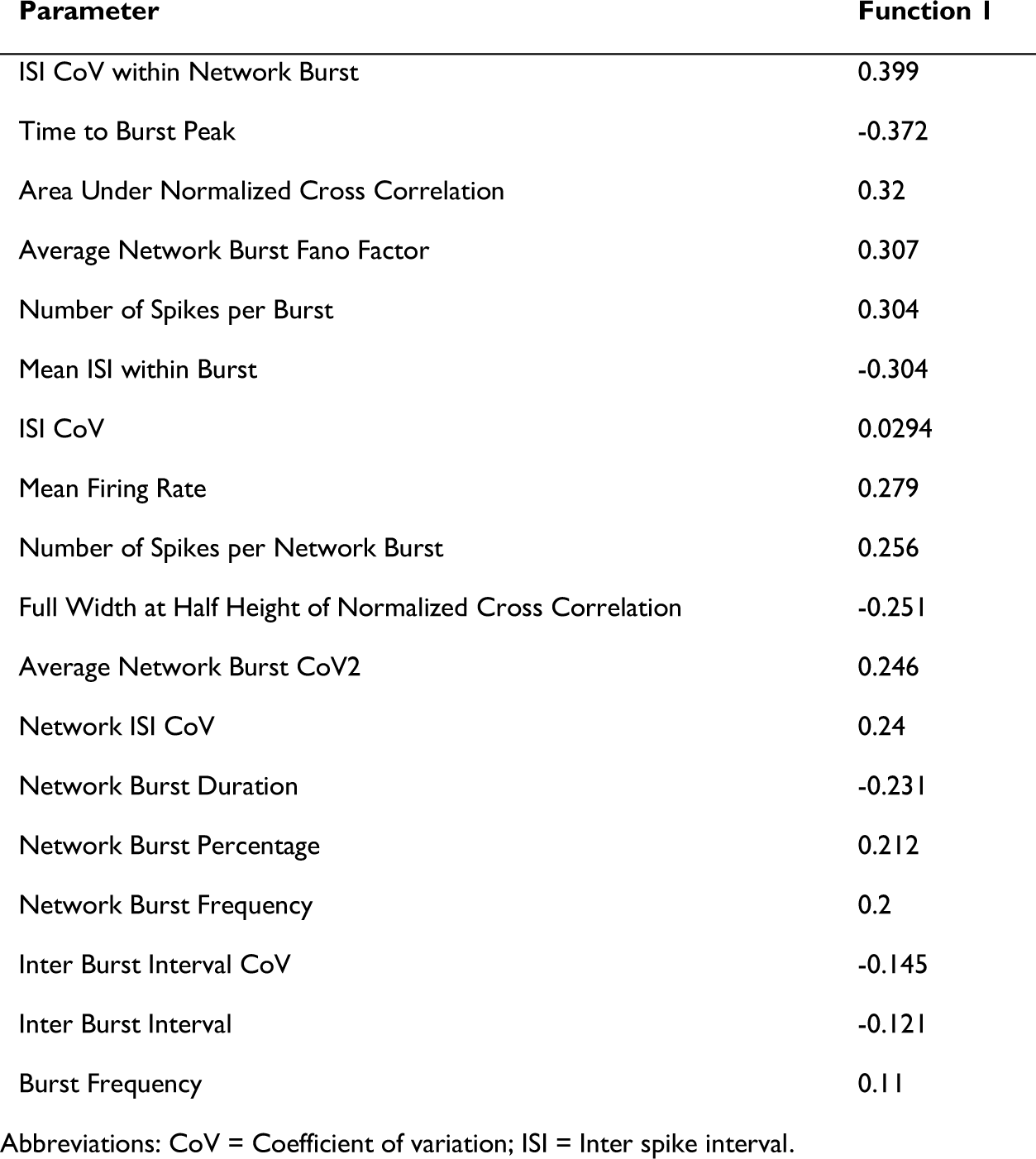
Structure matrix based on discriminant analysis of control and *CACNA1A*^+/−^ networks.

| Parameter | Function 1 |
| --- | --- |
| ISI CoV within Network Burst | 0.399 |
| Time to Burst Peak | -0.372 |
| Area Under Normalized Cross Correlation | 0.32 |
| Average Network Burst Fano Factor | 0.307 |
| Number of Spikes per Burst | 0.304 |
| Mean ISI within Burst | -0.304 |
| ISI CoV | 0.0294 |
| Mean Firing Rate | 0.279 |
| Number of Spikes per Network Burst | 0.256 |
| Full Width at Half Height of Normalized Cross Correlation | -0.251 |
| Average Network Burst CoV2 | 0.246 |
| Network ISI CoV | 0.24 |
| Network Burst Duration | -0.231 |
| Network Burst Percentage | 0.212 |
| Network Burst Frequency | 0.2 |
| Inter Burst Interval CoV | -0.145 |
| Inter Burst Interval | -0.121 |
| Burst Frequency | 0.11 |

Abbreviations: CoV = Coefficient of variation; ISI = Inter spike interval.

### Generation of rtTA-*Ngn2*-positive iPSCs and neuronal differentiation

In order to differentiate iPSCs into glutamatergic neurons, an inducible *Ngn2* gene cassette was introduced in iPSCs according to a previously published protocol.^19,22^ iPSCs were transduced with lentiviral vectors to stably integrate rtTA (pLV-EF1α>Tet3G:IRES:Neo) and *Ngn2* (pLV[TetOn]-Puro-TRE3G>mNeurog2([NM_009718.3])) transgenes into their genome. 48 hr after transduction, cells that carry both vectors were selected by applying G418 (25 μg/ml; Sigma-Aldrich, #G8168) and puromycin (0.5 µg/ml; Sigma-Aldrich, #P9620), of which concentrations were increased over time. The iPSCs that survived the selection process, were cultured in Essential 8^TM^ Flex Basal medium (Gibco) supplemented with primocin (Invivogen, #ant-pm-2), G418 (50 μg/ml) and puromycin (0.5 µg/ml) on Geltrex-(Gibco) coated plates at 37°C/5% CO_2_. Cells were passaged 1-2 times per week with ReLeSR (STEMCELL Technologies) when they reached 80-90% confluence. iPSCs were differentiated into cortical glutamatergic neurons by doxycycline-inducible *Ngn2* overexpression,^22^ according to a previously published protocol (Supplementary Methods and materials).^19^

### Micro-electrode array recordings and analysis

48-well Cytoview MEAs were used to record spontaneous neuronal network activity for 5 min in a Maestro Pro MEA system equipped with AxIS Navigator software (Axion BioSystems, Atlanta, GA, USA) at 37 °C and 5% CO_2_. Each well contains 16 electrodes in a 4×4 grid (diameter 50 µm, spacing 350 µm). The sampling frequency was set at 12.5 kHz. For spike detection, an adaptive threshold of ±6 standard deviations of the noise was applied. Spike, burst, and network burst parameters were extracted with AxIS Neural Metrics Tool software (Axion BioSystems). Electrodes were considered active when detecting at least 5 spikes per minute. Spikes were considered part of a burst when 5 spikes were detected with a maximum inter-spike interval (ISI) of 100 ms. Network bursts were determined with the built-in envelope algorithm with the threshold factor set at 2, minimum ISI at 100 ms, minimum 50% of electrodes active, and burst inclusion at 50%. For detection of network bursts during NMDA and AMPA receptor blockage, the ISI threshold algorithm was used with a minimum number of spikes per network burst of 500, a maximum ISI of 10 ms, and a minimum of 50% of electrodes active. Linear discriminant analysis was performed on data of eighteen parameters using SPSS (version 29.0.0, IBM Corporation, Armonk, NY, USA).

### In silico modeling

To investigate pathophysiological mechanisms underlying the spontaneous neuronal network activity, we employed a previously developed *in silico* model.^23^ In short, the model contains 100 Hodgkin-Huxley type neurons with voltage-gated sodium and potassium currents, and a slow, spike-dependent afterhyperpolarization current (sAHP). The neurons are sparsely connected through AMPA and NMDA receptors, where the properties of short-term synaptic depression (STD) were implemented. Virtual electrodes are used to mimic the MEA signal. We employed a recently introduced machine learning tool for simulation-based inference (sbi)^24^ to estimate the set of seven model parameters consistent with our experimental data. These parameters describe the most important cellular and synaptic mechanisms in the model (Supplementary Table 1). We generated 100.000 simulations with different parameter values. Both simulations and experimental measurements were analyzed in the same way. Spikes were detected using a threshold-based peak detection method, and the mean firing rate feature was calculated. Network bursts were detected using a firing rate threshold method as described earlier.^23^ From this, the network burst frequency, network burst duration, and network burst percentage features were calculated. The correlation coefficients between the binary spike train signals of all the electrodes were calculated to define the average correlation and standard deviation of the correlation features. To simulate control and *CACNA1A*^+/−^ networks, we used parameter values of the mode of the posterior distributions calculated with sbi. Simulations and experimental MEA data were compared using eight data features.

### Single cell electrophysiology

Single cell recordings in whole cell patch clamp configuration were performed at DIV42 as previously described.^25,26^ Coverslips were placed in a recording chamber on the stage of an Olympus BX51WI upright microscope (Olympus Life Science), equipped with infrared differential interference contrast optics, an Olympus LUMPlanFL N 60x water-immersion objective (Olympus Life Science), and a kappa MXC 200 camera system (Kappa optronics GmbH) for visualization. The recording chamber was continuously perfused with oxygenated (95% O_2_/5% CO_2_) artificial cerebrospinal fluid (aCSF) at 32°C containing (in mM): 124 NaCl, 1.25 NaH_2_PO_4_, 3 KCl, 26 NaHCO_3_, 11 Glucose, 2 CaCl_2_, 1 MgCl_2_. Patch pipettes were pulled from borosilicate glass with filament and fire-polished ends (ID 0.86 mm, OD1.05 mm, resistance 5–8 MΩ, Science Products GmbH) using the Narishige PC-10 micropipette puller. For recordings of intrinsic properties in current clamp mode and spontaneous excitatory postsynaptic currents (sEPSCs) in voltage clamp mode, pipettes were filled with a potassium-based solution containing (in mM): 130 K-Gluconate, 5 KCl, 10 HEPES, 2.5 MgCl_2_, 4 Na_2_-ATP, 0.4 Na_2_-GTP, 10 Na_2_-phosphocreatine, 0.6 EGTA (with pH adjusted to 7.2 and osmolarity to 290 mOsmol). All recordings were acquired using a Digidata 1140A digitizer and a Multiclamp 700B amplifier (Molecular Devices), with a sampling rate at 20 kHz and a lowpass filter at 1kHz. Recordings were not corrected for liquid junction potential (−15.646 mV; LJPcalc software, https://swharden.com/LJPcalc).^27^ Recordings were not analyzed if series resistance was above 25 MΩ or when it became higher than 10% of the membrane resistance. Resting membrane potential (Vrmp) was determined directly after reaching whole-cell configuration. Further analysis of active and passive membrane properties was conducted at a holding potential of −60 mV. Active intrinsic properties were measured with a stepwise current injection protocol ranging from −30 pA to +70 pA. sEPSCs were recorded for 10 min continuously at a holding potential of −60 mV. Burst rate, duration, and amplitude of bursting activities detected in sEPSC recordings were analyzed using a custom-made code developed in MATLAB (The Mathworks, Natick, MA, USA). Amplitude and frequency of individual events within sEPSC recordings were not analyzed, as there were too many interruptions by bursts. Action potential intrinsic properties were analyzed with the Action Potential Search algorithm of Clampfit 11.2 (Molecular devices). We assessed the properties of every first elicited action potential. Where applicable, measurements are reported relative to the threshold of each action potential.

### RNA-sequencing

Neuronal cultures of three replicates per cell line were harvested at DIV49. RNA was isolated with the Quick-RNA Microprep kit (Zymo Research) according to manufacturer’s instructions. The quality of the RNA was checked using Agilent’s Tapestation system (RNA High Sensitivity ScreenTape and Reagents). RNA Integrity Number values ranged between 8.6 and 9.1. cDNA libraries were generated with iScript (Bio-Rad, #1708890), and sequenced with a paired-end read length on an Illumina NovaSeq 6000 platform at GenomeScan B.V. Leiden. The average number of reads was >40 million. Reads were mapped to the human GRCh38.p13 (Homo_sapiens.GRCh38.dna.primary_assembly.fa) reference genome with STAR2 v2.7.10 and sorted with samtools v1.10 and counted with HTSeq v0.11.0. Raw count matrices were loaded in R v4.2.1.

### RNA-sequencing data analyses

To handle Ensemble IDs associated with identical gene symbols, we chose the IDs exhibiting the most elevated expression for each sample. Genes exhibiting minimal expression (less than 5 normalized counts, using DESeq2^28^ size factors method, in at least 3 samples were excluded. The remaining counts were normalized using voom function from the limma R package.^29^ Principal component analysis (PCA) was performed on scaled data, with the prcomp function.

Differential expression analysis between *CACNA1A*^+/−^ and control neurons was performed with limma. First, the count matrix was converted into a DGEList using DGEList function from edgeR.^30^ Voom normalization was then applied, taking into account normalization factors determined by edgeR’s calcNormFactors function and factoring in the condition (*CACNA1A*^+/−^ or control). To address within-cell line variability, the duplicateCorrelation function from limma was applied. Finally, limma’s functions lmFit, makeContrasts, contrasts.fit, and eBayes were used to derive the condition coefficient and its corresponding p-value for each gene. Genes were identified as differentially expressed if they had a Benjamini-Hochberg (BH)-adjusted p-value below 0.05 and a Log2 fold change greater than 0.58. Hierarchical clustering of samples, focusing on differentially expressed genes (DEGs), was conducted using the heatmap.2 function from the gplots v3.1.3 R package. This analysis utilized the Euclidean distance metric and employed the “ward.D2” clustering method. Enrichment analysis was separately conducted on down-regulated and up-regulated DEGs, using the go function from the R package gprofiler2 v0.2.1. To manage redundancy among enriched gene ontology terms (including biological processes, cellular components, or molecular functions), we performed clustering analysis and aggregated terms with high semantic similarity, utilizing the functions calculateSimMatrix and reduceSimMatrix with a threshold of 0.9 from the rrvgo v1.2.0 R package. For heatmaps depicting the expression patterns of synaptic genes across the samples, the normalized counts were scaled between samples. All plots were generated using custom code based on ggplot2 v.3.4.0 functions in R.

### Immunocytochemistry

For neuronal morphological reconstruction and quantification of synapse density, neurons were fixated at DIV42 with 4% paraformaldehyde/4% sucrose for 15 min and permeabilized with 0.2% Triton X-100 (Sigma Aldrich) in PBS for 10 min at room temperature. The cells were blocked in blocking buffer consisting of 5% normal goat serum (Invitrogen) in PBS for 1 hr at room temperature. Primary antibodies (guinea pig anti-MAP2 (1:1000; Synaptic Systems, #188 004), rabbit anti-Synapsin1 (1:500; Sigma-Aldrich, #AB1543P), mouse anit-Homer1 (1:500; Synaptic Systems, #160 011)) were diluted in blocking buffer and incubated overnight at 4°C. Secondary antibodies (goat anti-guinea pig Alexa Fluor 647 (1:1000; Invitrogen), goat anti-rabbit Alexa Fluor 568 (1:1000; Invitrogen), goat anti-mouse Alexa Fluor 488 (1:1000, Invitrogen)) were incubated for 1 hr at room temperature. Cells were mounted with fluorescence mounting medium (DAKO). Images were captured by a Zeiss Axio Imager Z1.

### Neuronal morphological reconstruction

Somatodendritic reconstructions were performed using NeuroLucida 360 software (Version 11, MBF–Bioscience). For somatodendritic analyses, only fully visible neurons with at least two dendrites and non-overlapping somas were considered for further analysis. We quantified the total number of primary dendrites, number of dendritic ends and nodes, the mean dendritic length, and the total dendritic length using NeuroLucida Explorer software. The mean dendritic length is calculated as total dendritic length divided by the total amount of primary dendrites. Furthermore, the complexity of the dendrites in a distance dependent manner was investigated by Sholl analysis using NeuroLucida Explorer. For each distance interval of 10 µm, the total dendritic length was quantified.

### Compounds

All drugs were prepared into concentrated stocks in ultrapure water, unless mentioned otherwise, and stored at −20 °C. Control networks were treated with 200 nM ω-agatoxin IVA (Sigma-Aldrich, #A6719) at DIV51 and recorded on MEA for 30 minutes after the compound was diluted into the medium. Both control and *CACNA1A*^+/−^ networks at DIV49 were treated acutely with 100 µM D-2-amino-5-phosphonovalerate (D-AP5; Tocris, #0106) for 60 min, after which 10 µM 1-Naphthyl acetyl spermine trihydrochloride (Naspm; Tocris, #2766) was added for 20 min. All networks received a 20-min 50 µM 2,3-Dioxo-6-nitro-1,2,3,4-tetrahydrobenzo[f]quinoxaline-7-sulfonamide (NBQX, in DMSO; Tocris, #0373) treatment on top of that. Neuronal network activity was recorded on MEA during the full treatment period. *CACNA1A*^+/−^ networks on MEA at DIV51 were treated with 20 µM 4-aminopyridine (4-AP; Tocris, #504-24-5) and recorded for 30 minutes after the compound was added. At single-cell level, *CACNA1A*^+/−^ neurons were treated with 20 µM 4-AP by bath application at DIV42.

### Statistical analysis

Statistical analysis was performed using GraphPad PRISM 9.0.0 (GraphPad Software, Inc., CA, USA). Data is shown in violin plots with dashed lines representing the median and dotted lines representing the quartiles, or in scatter or line plots with error bars expressing the mean ± standard error of the mean (SEM). In all figures, p-values are indicated as follows: <0.05 (*), <0.01 (**), <0.001 (***), <0.0001 (****). In each figure legend, the *n* indicates the number of wells or cells/independent differentiations. Normality was ensured using a Shapiro-Wilk test. Unless mentioned otherwise, comparison of two groups with non-Gaussian distributed data was performed with a Mann-Whitney test. If applicable, a post-hoc Bonferroni correction for multiple testing was performed on dependent parameters. For the line plots (Fig. 3F, 4B, 5E), two-way ANOVA with post hoc Bonferroni correction was performed. When comparing three or more groups (Fig. 7C, E) a one-way ANOVA with Bonferroni correction for multiple testing (for normally distributed data) or Kruskal-Wallis test with Dunn’s correction for multiple testing (for non-Gaussian distributed data) was performed.

### Data availability

All data and codes generated in this study are available from the corresponding author upon request. This study did not generate any novel reagents apart from iPSC lines. The iPSC lines used in this study are available upon request with a completed Materials Transfer Agreement from the corresponding author.

## Results

### *CACNA1A* haploinsufficiency leads to altered network synchronization in excitatory neuronal networks

In order to assess how loss-of-function of Ca_v_2.1 influences neuronal network activity, we first investigated whether acute blockage of Ca_v_2.1 affects spontaneous firing patterns of excitatory neuronal networks. To this end, we differentiated a control line (UCSFi001-A) to excitatory neurons via forced *Ngn2* overexpression (Fig. 1B).^19^ Upon seven weeks of network maturation, we applied the specific Ca_v_2.1 blocker, ω-agatoxin IVA,^31^ and recorded network activity with MEAs. Untreated wells showed the well-characterized and benchmarked network activity pattern of control networks, consisting of spikes as well as bursts and synchronous network bursts (Supplementary Fig. 2A), the latter of which reflects synchronization of the network activity.^32^ In sharp contrast, wells treated with ω-agatoxin IVA showed decreased global activity, reflected by a lower mean firing rate, with almost complete desynchronization of the network (i.e. lower network burst frequency and network burst percentage) (Supplementary Fig. 2A-B).

Thus, complete and acute pharmacological block of Ca_v_2.1 results in substantial reduction of network synchronization. To examine the effect of *CACNA1A* haploinsufficiency, we generated two heterozygous iPSC clones (*CACNA1A*^+/−^ (1) and *CACNA1A*^+/−^ (2)) with a frameshift mutation in exon 8 of *CACNA1A* via CRISPR/Cas9 in the same control line (Fig. 1A, Supplementary Fig. 1A).^18^ RT-qPCR confirmed reduced expression of *CACNA1A* (Supplementary Fig. 1F) and Western blot analysis showed, although not significant, a reduced protein level of CACNA1A in both *CACNA1A*^+/−^ lines (Supplementary Fig. 1H). In order to ensure comparable properties for network formation, we verified that cell densities between the different neuronal cultures were equal upon three weeks of differentiation (Supplementary Fig. 1I).

We next compared network activity of control and *CACNA1A*^+/−^ neurons (Fig. 1A-C). In contrast to a full and acute block of Ca_v_2.1 by ω-agatoxin IVA, *CACNA1A* haploinsufficiency did not lead to a substantial reduction in network synchronization, but to altered network synchronization (Fig. 1D). To uncover the network parameters that best distinguished the control and *CACNA1A*^+/−^ networks, we performed linear discriminant analysis^33,34^ on eighteen MEA parameters (Fig. 1E). This analysis unveiled the inter-spike interval (ISI) coefficient of variation (CoV) within network burst and the time to burst peak as the main discriminating parameters (Table 1). Specifically, we detected a lower ISI CoV within network burst and longer time to burst peak for *CACNA1A*^+/−^ networks (Fig. 1F). Thus, the synchronicity within network bursts was altered. Characterization of the networks by multiple MEA parameters describing general activity and bursting behavior,^32^ showed that the mean firing rate remained unaltered, whereas the network burst frequency and network burst percentage were lowered in *CACNA1A*^+/−^ networks (Fig. 1G). The network burst duration in these networks was higher than in control networks (Fig. 1G). These results show that there is a developmental and/or dosage effect of Ca_v_2.1 on the network burst activity. In conclusion, *CACNA1A* haploinsufficiency leads to an altered spontaneous neuronal network activity pattern.

### *In silico* modeling of control and *CACNA1A*^+/−^ networks reveal potential mechanisms driving the network phenotype

To investigate which pathophysiological mechanisms may cause the aberrant neuronal network activity, we employed our recently developed *in silico* model that has been proven to identify mechanisms underlying the MEA phenotype in Dravet syndrome.^23^ We simulated *in silico* the activity that would be exhibited by control and *CACNA1A*^+/−^ neuronal networks grown on MEA (Fig. 2A). We identified the most probable combinations of mechanistic parameters leading to the correct modeling of the empirical *in vitro CACNA1A*^+/−^ phenotype and compared these to the parameter values of control simulations (Fig. 2B, Supplementary Fig. 3). We found that short term depression (STD) of excitatory glutamatergic synaptic signaling remained unchanged. In contrast, a lower conductance of the sodium and potassium channels together with impaired excitatory synaptic function (i.e., either lower synaptic strengths, a lower connection probability, or a higher NMDA/AMPA ratio), were needed to simulate the *CACNA1A*^+/−^ network activity. We found that the *in silico* model adequately recapitulated the activity recorded in the *in vitro* control and *CACNA1A*^+/−^ neuronal networks (Fig. 2C). These results suggest that reduced sodium and potassium currents, as well as reduced synaptic function, might drive the *CACNA1A*^+/−^ network phenotype.

**Figure 2.**
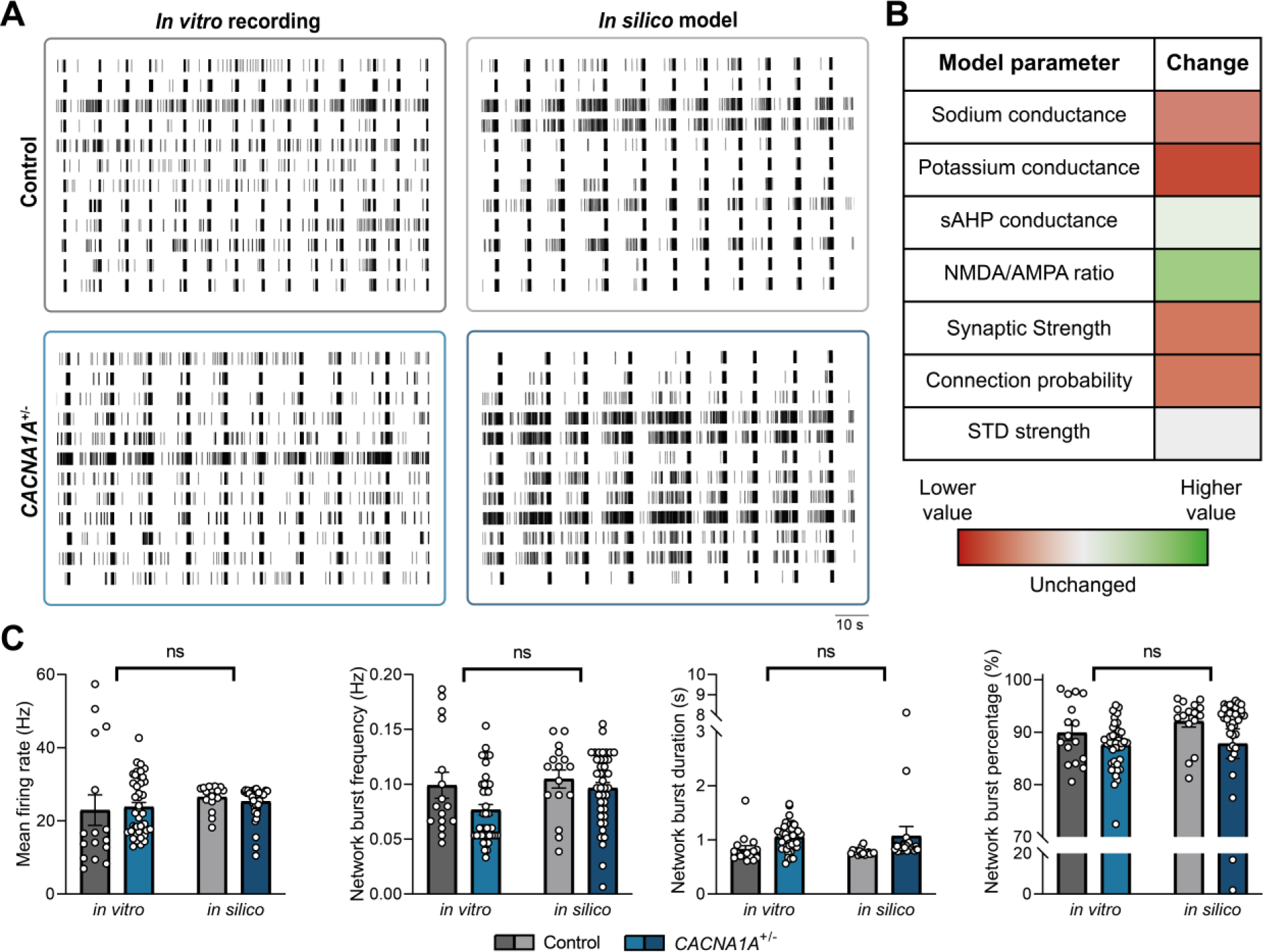
*In silico* model of *CACNA1A*^+/−^ networks replicates *in vitro* phenotype by altering synaptic function and sodium and potassium channel conductance. (**A**) Representative 2-min rasterplots of spontaneous activity from an *in vitro* control and *CACNA1A*^+/−^ network (left panel) and simulated activity from an *in silico* control and *CACNA1A*^+/−^ model (right panel). (**B**) Heatmap showing the magnitude (color intensity) and direction (red or green) of change for each of the *in silico* model parameters to transition from the control model to the *CACNA1A*^+/−^ model. (**C**) Quantification of network parameters including mean firing rate, network burst frequency, network burst duration and network burst percentage. *n = 16/3* for Control for both *in vitro* and *in silico*; *n = 41/3* for *CACNA1A*^+/−^ ((1): *21/3* and (2): *21/3*). for both *in vitro* and *in silico*. Data represent means ± SEM, ns *p > 0.05,* Means were compared with a two-way ANOVA with Bonferroni correction for multiple testing.

### *CACNA1A*^+/−^ neurons show synaptic deficits with augmented contribution of GluA2-lacking AMPA receptors

Next, we tested the *in silico* predicted altered synaptic function in *CACNA1A*^+/−^ neurons in the *in vitro* networks. As mentioned above, reduced connection probability, reduced synaptic strength, or a higher NMDA/AMPA ratio could all explain the *CACNA1A*^+/−^ network phenotype. To this end, we first measured synapse density (Fig. 3A). Both the number of Synapsin1/2 and Homer1 puncta were decreased in *CACNA1A*^+/−^ neurons, as well as their colocalization (Fig. 3B), indicating a reduced number of functional synapses. Somatodendritic structure and synapse formation of neurons are highly correlated, so a reduced number of synapses may affect the size of the dendritic arbor.^35^ To investigate this, we reconstructed control and *CACNA1A*^+/−^ neurons to study their somatodendritic structure (Fig. 3C). Overall, we did not observe very prominent differences between both genotypes on the qualitative level. However, detailed morphometrical quantification of revealed an increased soma size for *CACNA1A*^+/−^ neurons (Fig. 3D). Even though the number of primary dendrites and dendritic ends, as well as the total dendritic length showed no significant differences (Fig. 3D), Sholl analysis revealed a significant increase in dendritic length close to the soma, thus, increased perisomatic dendritic complexity (Fig. 3E-F).

**Figure 3.**
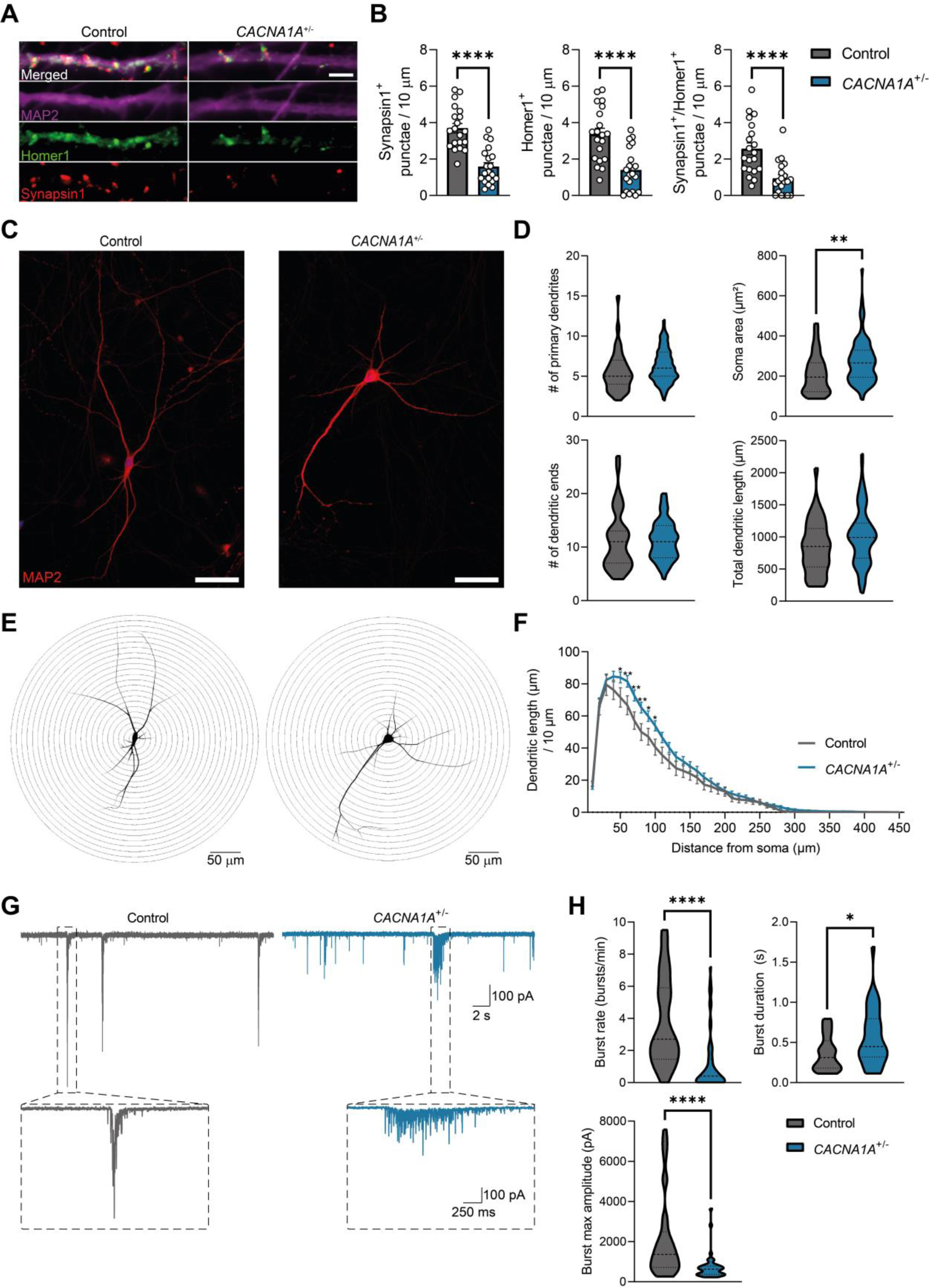
*CACNA1A*^+/−^ neurons show reduced synaptic density, altered somatodendritic morphology and reduced synaptic function. (**A**) Representative images of control and *CACNA1A*^+/−^ dendrites at DIV42 (63x magnification) stained with microtubule-associated protein 2 (MAP2), Homer1 and Synapsin1 (scale bar 5 μm). (**B**) Quantification of the number of Synapsin1, Homer1 and co-localized Synapsin1/Homer1 punctae per 10 μm MAP2-positive dendritic length. *n = 20/3* for Control; *n = 20/3* for *CACNA1A*^+/−^ ((1): *10/3* and (2): *10/3*). Data represent means ± SEM, with individual datapoints representing the mean number of punctae per neuron. *\*\*\*\*p < 0.0001*, Mann–Whitney test. (**C**) Representative images of control and *CACNA1A*^+/−^ neurons at DIV42 (20x magnification) stained with MAP2 (scale bar 50 μm). (**D**) Quantification of somatodendritic parameters including number of primary dendrites, soma area, number of dendritic ends and total dendritic length. *n = 39/2* for Control; *n = 92/2* for *CACNA1A*^+/−^ ((1): *45/2* and (2): *47/2*). Dashed line represents the median, dotted line represents the quartiles. *\*\*p < 0.01,* Mann–Whitney test. (**E**) Representative somatodendritic reconstruction of control and *CACNA1A*^+/−^ neurons at DIV42 with 10-μm Sholl rings placed from the center soma outward. (**F**) Quantification per 10-μm Sholl ring of the total dendritic length per ring. *N = 39/2* for Control; *n = 91/2* for *CACNA1A*^+/−^ ((1): *44/2* and (2): *47/2*). Data represent means ± SEM. \**p < 0.05*, \*\**p < 0.01*, two-way ANOVA with post hoc Bonferroni correction. (**G**) Representative 30-sec whole-cell voltage-clamp recordings of spontaneous excitatory postsynaptic currents (sEPSCs) of control and *CACNA1A*^+/−^ neurons at DIV42, with a 2.5-sec zoom-in of an sEPSC burst. (**H**) Quantification of burst parameters including burst rate, burst duration and burst max amplitude. *n = 37/8* for Control; *n = 57/8* for *CACNA1A*^+/−^ ((1): *34/7* and (2): *23/4*). Dashed line represents the median, dotted line represents the quartiles. \**p < 0.05*, *\*\*\*\*p < 0.0001,* Mann–Whitney test.

To uncover the net change in functional synapses, we performed functional analysis by recording spontaneous excitatory postsynaptic currents (sEPSCs) (Fig. 3G). Control neurons frequently showed closely clustered synaptic inputs, resembling the (network) bursts detected on the MEA. We detected a reduced burst rate and increased burst duration in *CACNA1A*^+/−^ neurons, resembling the output at the network level (Fig. 3H). The maximum amplitude of the bursts was decreased in *CACNA1A*^+/−^ neurons, indicating that these neurons could have a reduced synaptic strength (Fig. 3H).

Lastly, to investigate whether the contribution of different glutamate receptors is altered in *CACNA1A*^+/−^ networks as predicted by the *in silico* model, we pharmacologically blocked NMDA and AMPA receptors, and compared the effect on control and *CACNA1A*^+/−^ networks.^33^ Consistent with the literature, inhibition of NMDA receptors by D-AP5 had no significant effect on the network burst frequency in control cells (Fig. 4A-B). Similarly, D-AP5 did not affect *CACNA1A*^+/−^ network activity (Fig. 4A-B), suggesting that NMDA receptors do not play an important role in maintaining network bursts. Blockage of GluA2-lacking AMPA receptors by Naspm slightly decreased network burst frequency in control networks (Fig. 4A-B, Supplementary Fig. 2C-D), but interestingly showed a profound effect on the *CACNA1A*^+/−^ network (Fig. 4C). The effect of Naspm on control networks decreases over development, having no effect anymore at DIV49 (Supplementary Fig. 2C-E). This indicates that GluA2-lacking receptors are replaced by GluA2-containing receptors, a process previously described in rodent models.^36^ The profound effect of Naspm on the *CACNA1A*^+/−^ network might indicate that *CACNA1A*^+/−^ neurons are developmentally delayed in the transition of GluA2-lacking to GluA2-containing receptors. Treatment of both networks with NBQX, a generic inhibitor of AMPA receptors, showed that AMPA receptors are essential for network synchronization in both control and *CACNA1A*^+/−^ networks (Fig. 4A-B).

**Figure 4.**
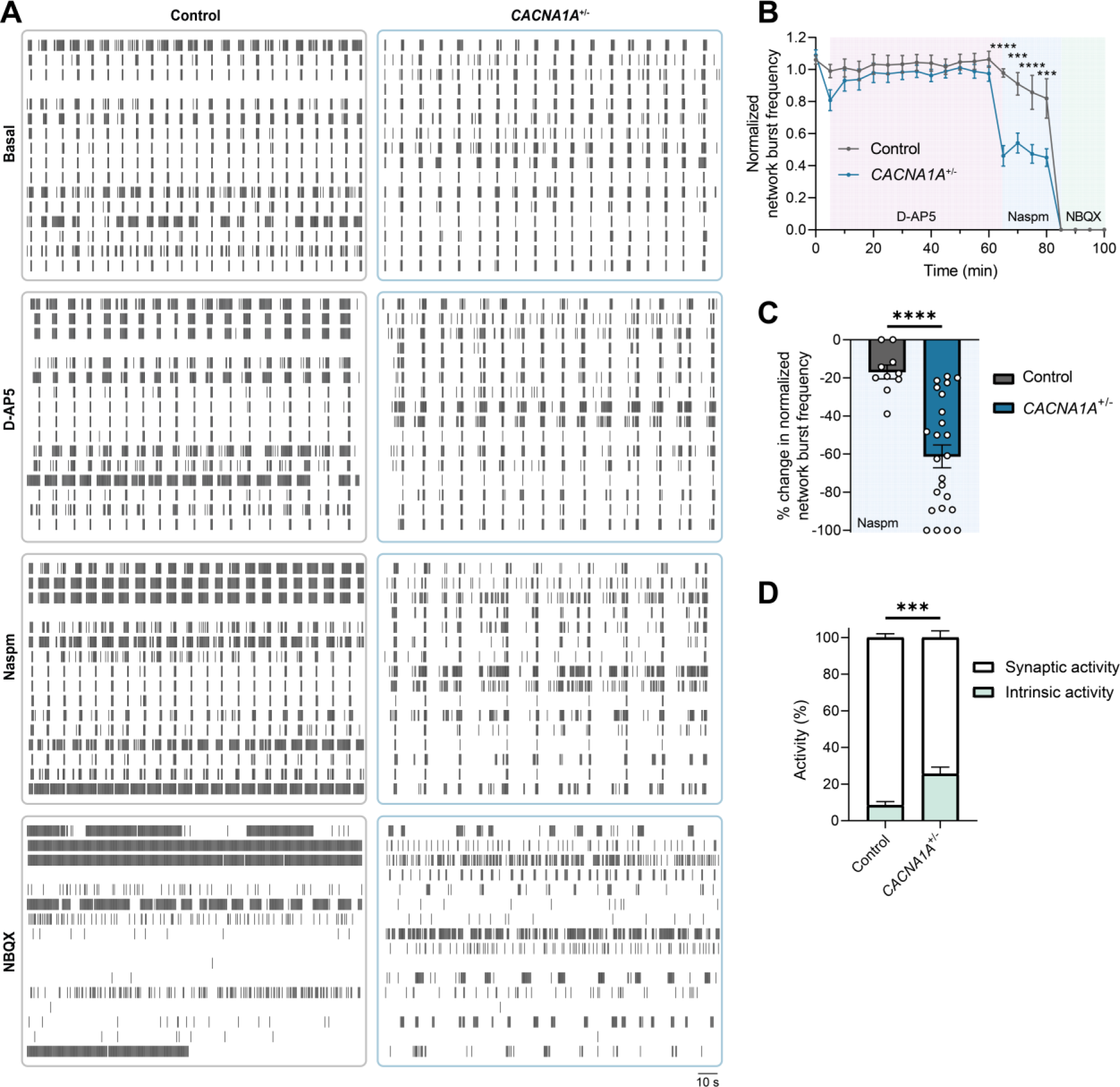
*CACNA1A*^+/−^ neurons have preserved GluA2-lacking AMPA receptor activity and show increased intrinsic activity. (**A**) Representative 3-min rasterplots of spontaneous activity from control and *CACNA1A*^+/−^ networks at DIV49. Where indicated, the cells were either non-treated (Basal) or treated with 100 μM D-AP5 to block NMDA receptors, 10 μM Naspm to block GluA2-lacking AMPA receptors, or 50 μM NBQX to block AMPA receptors. (B) Quantification of the normalized network burst frequency over time for control and *CACNA1A*^+/−^ networks. Networks were treated D-AP5 for 1 hr. After 1 hr, the GluA2-lacking AMPA receptor blocker Naspm was added for 20 min, after which NBQX was added for 20 min. The values are normalized to non-treated wells. (**C**) Quantification of change in normalized network burst frequency in the first 100 sec after adding Naspm for control and *CACNA1A*^+/−^ networks. The values are normalized to the activity 5 min before adding Naspm (when treatment of D-AP5 is ongoing) for each well. (**D**) Quantification of the percentage of intrinsic activity (mean firing rate after D-AP5, Naspm, and NBQX) normalized to the total activity (mean firing at time = 0 min) for each well. *n = 10/2* for Control; *n = 24/2* for *CACNA1A*^+/−^. ((1): *12/2* and (2): *12/2*). Data represent means ± SEM. *\*\*\*p < 0.001, ****p < 0.0001*, two-way ANOVA with post hoc Bonferroni correction for (**B**), Mann-Whitney test for **(C)** and (**D**).

Network activity eventually is a summation of intrinsically fired action potentials of individual neurons and synaptic communication between neurons. This intrinsically generated non-synaptically driven activity of neurons is on the one hand considered to be a contributor to network formation and maturation during development and, on the other hand, is also influenced by the amount of synaptic activity from the surrounding network. After establishing a complete suppression of glutamatergic synaptic communication by blocking NMDA as well as AMPA receptors, we yielded a network in which only non-synaptically driven spiking activity, henceforth referred to as intrinsic activity, of control and *CACNA1A*^+/−^ neurons was detectable via the MEA. In contrast to the synaptic deficits, quantification of the contribution of intrinsic activity to the total activity yielded an overactive intrinsic phenotype in *CACNA1A*^+/−^ neurons (Fig. 4D), suggesting that in *CACNA1A*^+/−^ networks, the individual neurons may show increased excitability.

To summarize, these combined results show that a decrease in synaptic density and synaptic strength, as well as preserved GluA2-lacking AMPA receptor contribution, and increased intrinsic activity underlie the neuronal network phenotype in our *CACNA1A*^+/−^ *in vitro* model.

### *CACNA1A*^+/−^ neurons display increased intrinsic excitability through reduced potassium channel function and expression

To corroborate our network observations at the single-cell level, we recorded intrinsic electrophysiological properties of control and *CACNA1A*^+/−^ neurons (Fig. 5A-E). We did not observe any significant changes in the passive properties of the cells (i.e. capacitance, membrane resistance, and the resting membrane potential (Vrmp) (Fig. 5D). However, when examining the active properties, we detected a lower rheobase (Fig. 5D), suggesting that, in line with the network data, individual *CACNA1A*^+/−^ neurons show increased intrinsic excitability. We also observed a longer action potential decay time and a reduced afterhyperpolarization (AHP) amplitude following the action potential peak (Fig. 5B-D). These results suggest a decreased conductance for voltage-gated and calcium-activated potassium channels, similar to the *in silico* model predictions. We found no significant difference in the action potential rise time (Fig. 5D), indicating that the action potential-related sodium conductance was not changed in *CACNA1A*^+/−^ neurons. To dissect how a lower conductance of both voltage-gated and calcium-activated potassium channels would influence the neurons’ tendency to fire action potentials, we constructed an input/output curve. We observed an increased number of action potentials per current injection in *CACNA1A*^+/−^ neurons (Fig. 5E), indicating that, consistent with a lower rheobase, individual *CACNA1A*^+/−^ neurons show increased intrinsic excitability.

**Figure 5.**
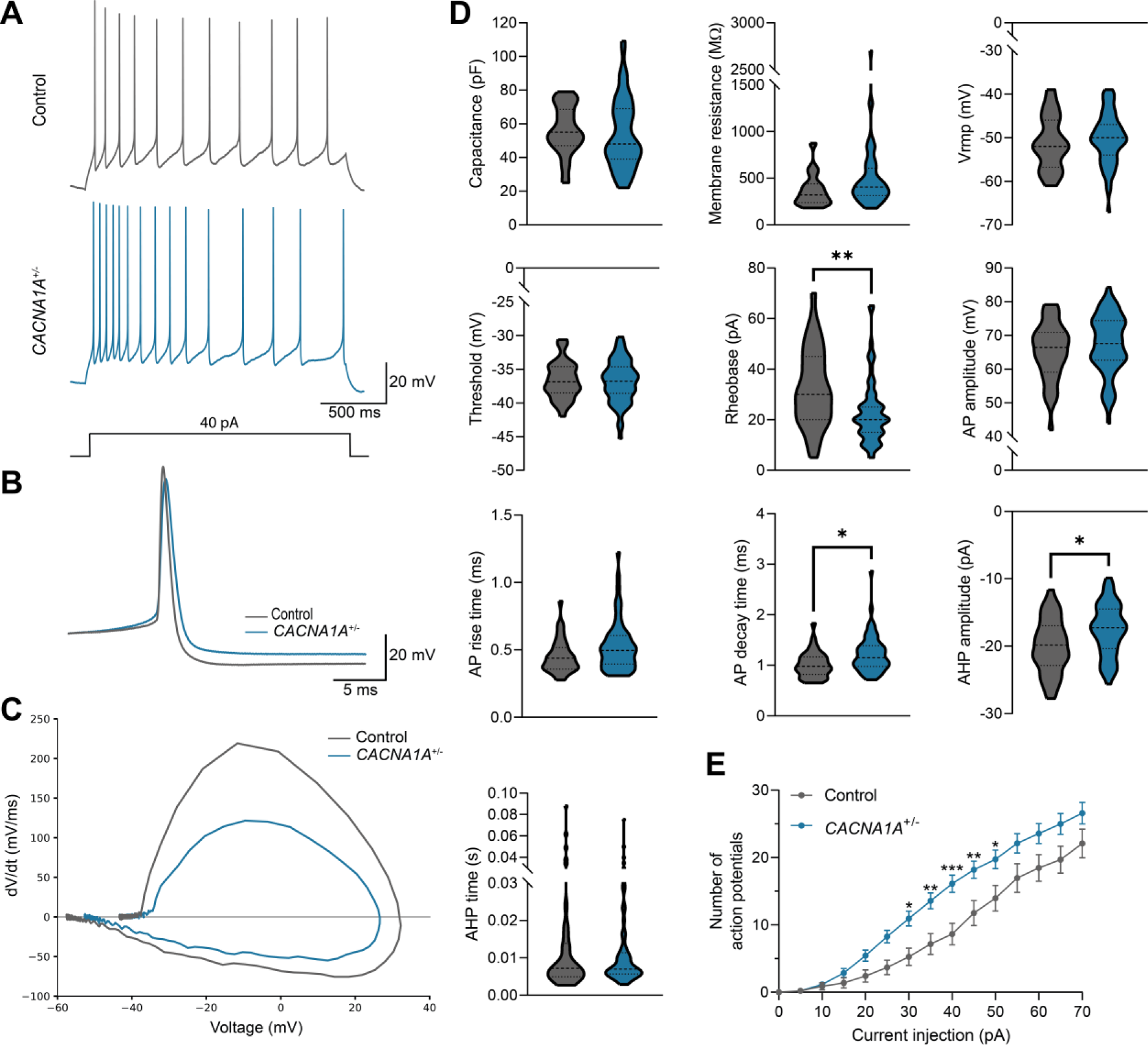
*CACNA1A*^+/−^ neurons display increased intrinsic excitability through reduced function of voltage-gated and calcium-activated potassium channels. (**A**) Representative 2.3-sec firing patterns of a control and *CACNA1A*^+/−^ neuron, recorded using a 2-sec pulse of 40 pA in whole-cell current clamp at DIV42. (**B**) Representative outline of single action potential waveforms. (**C**) Representative phase plot of the single action potential as plotted in (**B**). (**D**) Quantification of passive and active intrinsic properties as indicated. Dashed line represents the median, dotted line represents the quartiles. (**E**) Quantification of number of action potentials per 2-sec current injection. *n = 37/8* for Control; *n = 63/8* for *CACNA1A*^+/−^ ((1): *36/7* and (2): *27/4*). Data represent means ± SEM. \**p < 0.05*, *\*\*p < 0.01, ***p < 0.001*, Mann-Whitney test for (**D**), two-way ANOVA with post hoc Bonferroni correction for (**E**). Abbreviation: AP = action potential.

To investigate whether the decreased conductance for voltage-gated and calcium-activated potassium channels of *CACNA1A*^+/−^ neurons was also reflected by transcriptional changes, we performed RNA-sequencing (RNA-seq) of control and *CACNA1A*^+/−^ neurons. Principal component analysis (PCA) segregated control and *CACNA1A*^+/−^ neurons, indicating transcriptional changes in neuronal networks due to *CACNA1A* haploinsufficiency (Fig. 6A). We quantified the differentially expressed genes and identified 858 down-regulated and 929 up-regulated genes (Fig. 6B). The changes were consistent across all samples of the same genotype (Supplementary Fig. 4A). Gene ontology (GO) term analysis revealed the biological process (BP) term ‘system development’ to be the most strongly enriched in *CACNA1A*^+/−^ neurons (Fig. 6C), including members of the protocadherin family. Genes down-regulated by *CACNA1A* haploinsufficiency were involved in the cellular compartments (CC) ‘neuron projection’ and ‘synapse’, and in the biological processes ‘nervous system development’ and ‘trans-synaptic signaling’. Molecular function (MF) and KEGG pathway analysis revealed the ‘calcium signaling pathway’, ‘neurotransmitter receptor activity’, ‘calcium ion binding’, and ‘gated channel activity’ as important affected pathways. Considering that ‘gated channel activity’ was identified by GO term analysis and that our electrophysiological data showed reduced potassium channel function, we next calculated the fold change of differentially expressed ion channels. When focusing on voltage-gated and calcium-activated potassium channels, we identified *KCNH4* and *KCNN2* to be up-regulated, and *KCNC2, KCND1, KCNA4, KCNN1, KCNQ3,* and *KCND2* to be down-regulated (Fig. 6D, Supplementary Fig. 4B).

**Figure 6.**
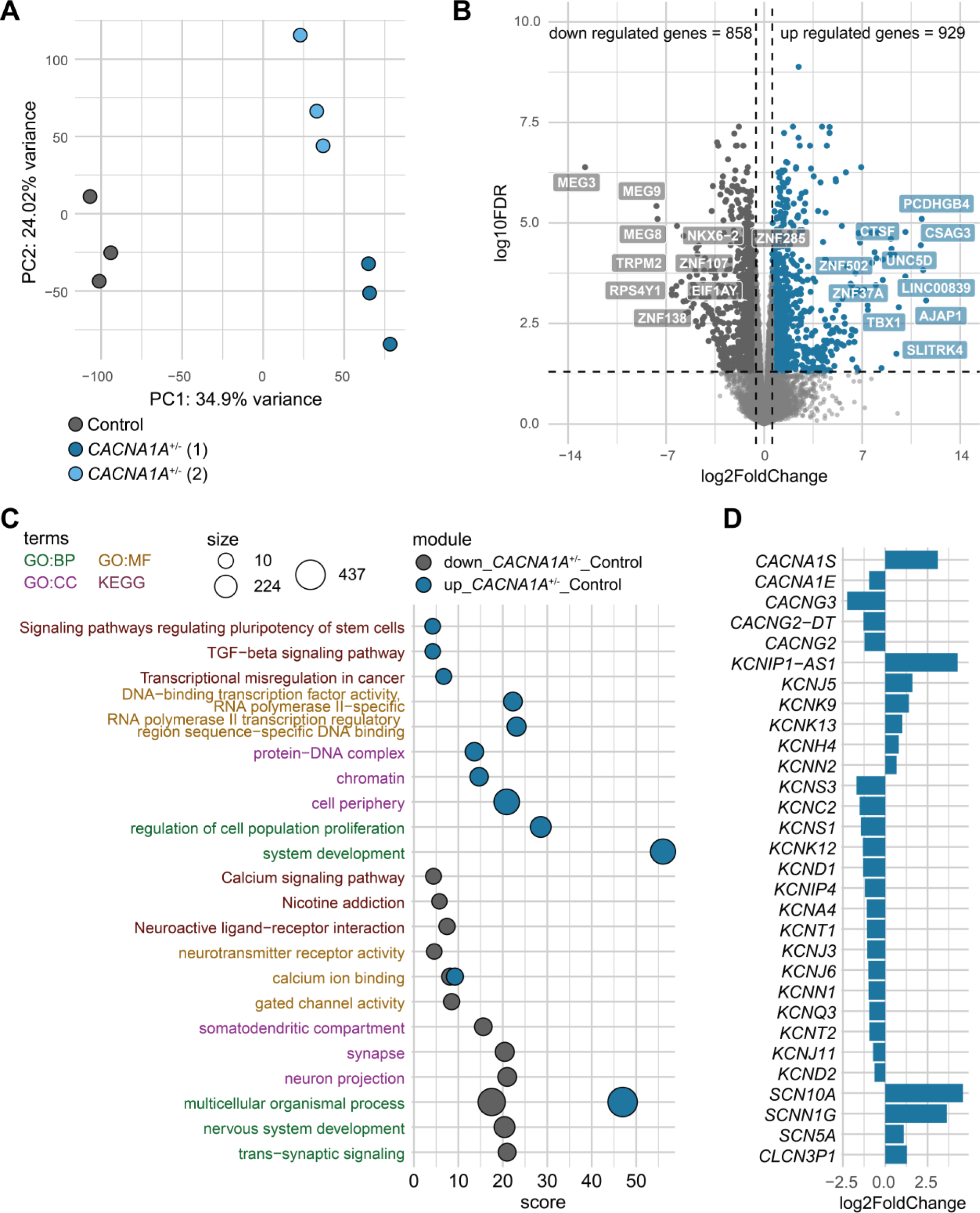
Potassium channel expression is altered in *CACNA1A*^+/−^ neurons. (**A**) Principal component analysis (PCA) of RNA-sequencing data of three control, three *CACNA1A*^+/−^ (1) and three *CACNA1A*^+/−^ (2) neuronal samples at DIV49. (**B**) Volcano plot depicting differential gene expression between control and *CACNA1A*^+/−^ neurons. Colored dots indicate differentially expressed genes (DEGs; absolute log_2_ fold change > 0.58 and adjusted p-value <0.05). Down-regulated genes in *CACNA1A*^+/−^ neurons are shown in grey, up-regulated genes are shown in blue. (**C**) Scatterplot of the enrichment analysis results depicting the top three significant gene ontology (GO) terms associated with up-regulated (blue dots) and down-regulated genes (grey dots) in *CACNA1A*^+/−^ neurons. Terms are ordered per ontological category (BP: biological process; CC: cellular component; MF: molecular function; KEGG: Kyoto Encyclopedia of Genes and Genomes). The size of the dots indicates the number of genes observed in the list of DEGs and the list of genes associated with the particular term. Score: negative log_10_ of the adjusted p-value resulting from the enrichment analysis. (**D**) Bar plot depicting the Log2FoldChange value of a selection of DEGs encoding ion channels. Fold change was calculated by normalizing gene expression level of *CACNA1A*^+/−^ neurons to control neurons.

In conclusion, these results suggest that *CACNA1A*^+/−^ neurons show reduced potassium channel function and expression, which may underlie the observed increased intrinsic excitability.

### Treatment of *CACNA1A*^+/−^ neurons with a potassium channel blocker partially rescues electrophysiological phenotype

Considering our data showing reduced potassium channel function and expression in *CACNA1A*^+/−^ neurons, it seems counterintuitive that *CACNA1A* patients with episodic ataxia benefit from treatment with 4-aminopyridine (4-AP), a non-selective voltage-gated potassium channel blocker.^37–42^ The original rationale of 4-AP treatment for episodic ataxia patients was to prolong the action potential to allow a higher calcium inflow into the cell.^37,41^ However, our recordings of *CACNA1A*^+/−^ neurons already showed such elongated action potentials. To investigate whether 4-AP could nevertheless rescue the network phenotype of *CACNA1A*^+/−^ neuron networks, we treated neurons with 4-AP for 30 min and recorded their network activity on MEA (Fig. 7A). Discriminant analysis based on the same parameters as in Fig. 1D showed a shift of the 4-AP-treated *CACNA1A*^+/−^ neuronal networks towards the control networks (Fig. 7B, Supplementary Table 2). This suggests that we partially rescued the *CACNA1A*^+/−^ network phenotype. Specifically, we detected no effect of 4-AP on the ISI CoV within network bursts, but the time to burst peak was restored to control levels (Fig. 7C). This raised the question about the effect of 4-AP on the intrinsic electrophysiological properties on single-cell level. At single-cell level, 4-AP indeed increased action potential decay time (Fig. 7D-E, Supplementary Fig. 5A-C), indicating successful blockage of voltage-gated potassium channels. AHP amplitude between all groups did not significantly change, but there was a trend towards a more negative AHP amplitude in *CACNA1A*^+/−^ neurons treated with 4-AP compared to untreated *CACNA1A*^+/−^ neurons (Fig. 7E). Previously, it has been shown that 4-AP enhances calcium-activated potassium channel currents in *CACNA1A*-deficient cells.^37,43^ In contrast, 4-AP treatment in control cells reduces the AHP amplitude (Supplementary Fig. 5A-C), consistent with the effect of 4-AP on rodent cortical interneurons.^44^ This shows that 4-AP has a differential effect in these two conditions. In conclusion, we observed that 4-AP partially rescues the electrophysiological phenotype of *CACNA1A*^+/−^ neurons, reflecting an effect restricted to the intrinsic properties of the neurons; the change in synaptic function cannot be rescued. In other words, only one of the two co-existing mechanisms that drive the electrophysiological phenotype of *CACNA1A*^+/−^ neurons can be influenced by the treatment of 4-AP.

**Figure 7.**
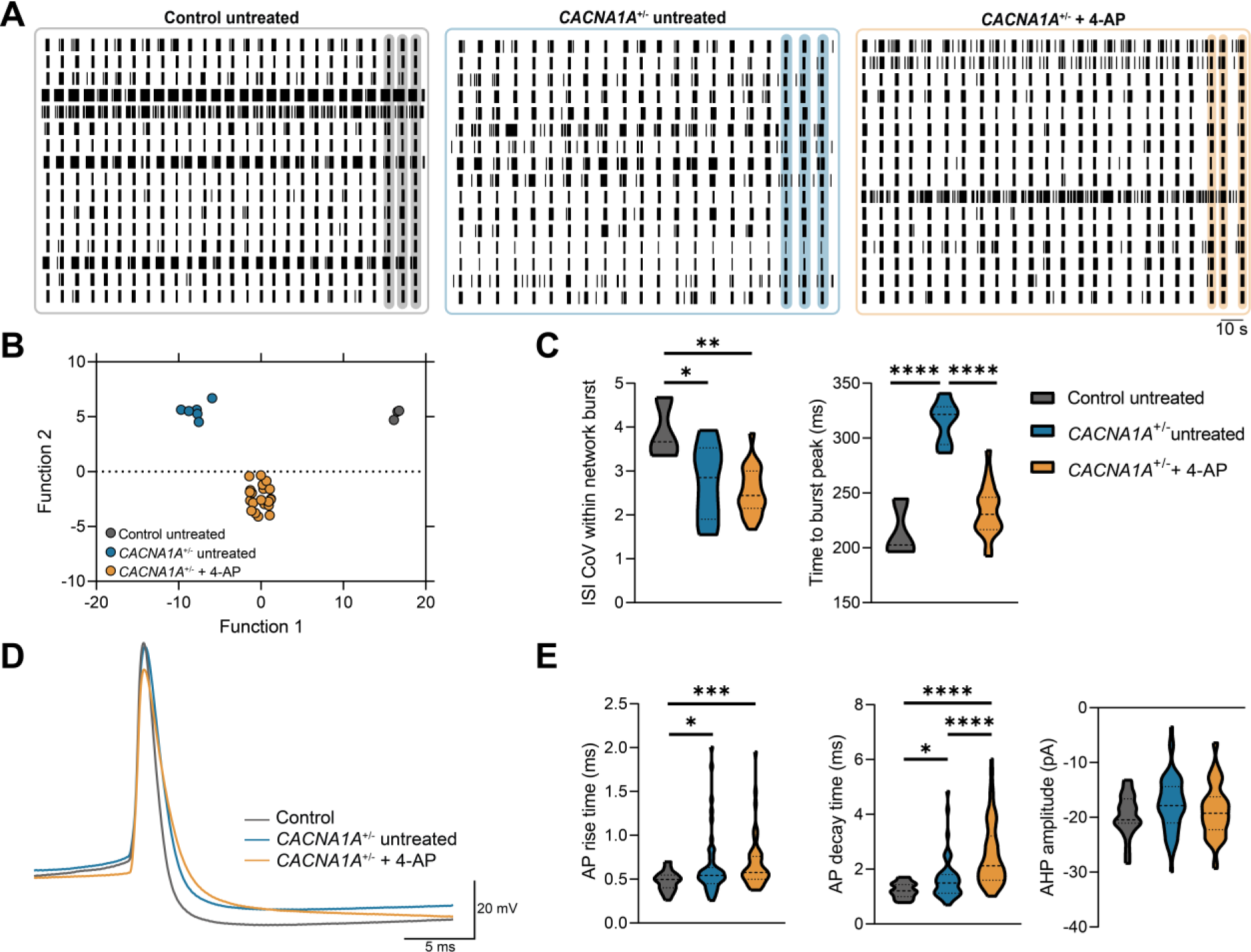
4-Aminopyridine (4-AP) partially rescues electrophysiological phenotype of *CACNA1A*^+/−^ neurons. (**A**) Representative 3-min rasterplots of spontaneous activity of control, *CACNA1A*^+/−^, and *CACNA1A*^+/−^ treated with 20 μM 4-AP for 30 min at DIV51. (**B**) Quantification of network parameters including inter spike interval (ISI) coefficient of variation (CoV) within network burst, and time to burst peak. *n = 3/2* for Control; *n = 6* for *CACNA1A*^+/−^ ((1): *3/2* and (2): *3/2*); *n = 22* for *CACNA1A*^+/−^ + 4-AP ((1): *11/2* and (2): *11/2*). Dashed line represents the median, dotted line represents the quartiles. \**p < 0.05, **p < 0.01, ****p < 0.0001*, one-way ANOVA with with post hoc Bonferroni correction. (**C**) Canonical score plots of control, *CACNA1A*^+/−^, and 4-AP-treated *CACNA1A*^+/−^ network based on discriminant analysis of eighteen MEA parameters as indicated in the structure matrix of Supplementary Table 2. (**D**) Representative outline of single action potential waveforms of control, *CACNA1A*^+/−^, and 4-AP-treated *CACNA1A*^+/−^ neurons at DIV42. (**E**) Quantification of active intrinsic properties as indicated. *n = 31/3* for Control; *n = 55/3* for *CACNA1A*^+/−^ ((1): *25/3* and (2): *30/3*); *n = 58* for *CACNA1A*^+/−^ + 4-AP ((1): *30/3* and (2): *28/3*). Dashed line represents the median, dotted line represents the quartiles. \**p < 0.05, ***p < 0.001, ****p < 0.0001*, Kruskal-Wallis test with post hoc Dunn’s correction. Abbreviation: AP = action potential.

## Discussion

In this study, we investigated pathological mechanisms underlying aberrant network function associated with *CACNA1A* haploinsufficiency using an iPSC-derived neuronal model. We differentiated iPSCs to glutamatergic neurons and first verified the validity of the use of glutamatergic neurons to model *CACNA1A* haploinsufficiency by treating control excitatory neuronal networks with ω-agatoxin IVA, which affected the spontaneous firing patterns. In mature excitatory neuronal networks with *CACNA1A* haploinsufficiency we observed an altered network synchronization.

To narrow down the possible cellular mechanisms that underlie the observed altered network synchronization, we employed a recently developed predictive and hypothesis generating *in silico* model.^23^ The cellular mechanisms identified by the *in silico CACNA1A*^+/−^ model were 1) decreased synaptic function with an increased NMDA/AMPA ratio, and 2) decreased sodium and potassium channel function. We tested and validated these findings in our *in vitro CACNA1A*^+/−^ model and indeed found a decrease in synaptic function with enhanced contribution of GluA2-lacking AMPA receptors, and increased intrinsic excitability through reduced potassium channel function. However, our empirical *in vitro* results were not entirely in line with the prediction of the *in silico* model, which suggested increased influence of NMDA receptors or decreased influence of AMPA receptors. This might be due to the fact that only GluA2-containing AMPA receptors were included in the *in silico* model. While the AMPA-current is modeled as a fast current that is instantaneous with the arrival of a presynaptic action potential, the NMDA-current is modeled as a slow current that can only be activated after prolonged stimulation (i.e., voltage-dependent). Since *in vitro* GluA2-lacking AMPA receptors are generally slower than GluA2-containing AMPA receptors and are blocked by polyamines in a voltage-dependent manner, they might be more similar to the modeled NMDA receptors than the GluA2-containing AMPA receptors. Thus, the prediction of increased NMDA receptors might reflect an increase in GluA2-lacking AMPA receptors.

In control neurons we generally observed a decreased contribution of GluA2-lacking receptors over development (Supplementary Fig. 2C, D), accompanied by a shortening of the network bursts (Supplementary Fig. 2E).^32^ The preserved contribution of GluA2-lacking receptors in *CACNA1A*^+/−^ neurons may explain the observed increased network burst duration (Fig. 1E), and may indicate that *CACNA1A*^+/−^ neurons are developmentally delayed. However, as the GluA2-lacking receptors are both sodium- and calcium-permeable,^45,46^ whereas GluA2-containing receptors are only sodium-permeable, it may also be a compensatory effect for *CACNA1A* haploinsufficiency.

Given the fact that Ca_v_2.1 plays an important role in synaptic communication,^47,48^ the observed reduced connection probability could be considered a direct effect of *CACNA1A* haploinsufficiency. As somatodendritic structure and synapse number are correlated, we expected less complex dendritic morphology of *CACNA1A*^+/−^ neurons, but instead found a more complex perisomatic dendritic organization. L-type voltage-gated calcium channels have been shown to be critical for dendritic growth and neuronal maturation in the developing cortex,^49–52^ suggesting that *CACNA1A* haploinsufficiency might lead to a lower complexity of the dendritic arborization. On the other hand, we identified up-regulation of many members of the protocadherin family, which play an important role in the development of dendrites, axons, and synapses.^53,54^ Future studies are needed to investigate how gene regulation of the protocadherin protein family is connected to *CACNA1A*.

We have characterized a synaptic deficit in *CACNA1A*^+/−^ neurons, which is in contrast to existing literature on a heterozygous rodent model.^55^ The synaptic deficit found in our study was surprisingly accompanied by an increased intrinsic excitability. In this study, we unprecedentedly exploited the inhibition of NMDA and AMPA receptors on MEAs to determine the fraction of intrinsic activity of the neurons in the network (Fig. 4D). Counterintuitively, *CACNA1A*^+/−^ neurons showed increased intrinsic activity, which was corroborated at the single-cell level (Fig. 5). We have recently shown that iPSC-derived neuronal networks undergo homeostatic plasticity,^56^ which indicates that these networks continuously strive for a certain level of activity. Hence, reduced synaptic function in *CACNA1A*^+/−^ neurons might trigger increased intrinsic activity. However, the flow of calcium through P/Q-type channels has also directly been linked to the recruitment of calcium-activated potassium channels in Purkinje neurons,^57,58^ which indicates that increased intrinsic excitability may also be a direct effect of *CACNA1A* haploinsufficiency. Remarkably, we identified the brain-specific *KCNN1* to be down-regulated in *CACNA1A*^+/−^ neurons, showing that this functional link is reflected at the transcriptional level. Reduced levels or activity of small conductance calcium-activated potassium channels have been associated with epilepsy and ataxia,^59,60^ two of the core phenotypes that are associated with *CACNA1A* haploinsufficiency. We also identified that *KCNN2* was up-regulated in *CACNA1A*^+/−^ neurons. Overexpression of this gene has been shown to modulate synaptic plasticity in rodent hippocampal neurons.^61^ The effect of *CACNA1A* haploinsufficiency and the concurrent up-regulation of *KCNN2* on synaptic plasticity remains to be investigated.

In summary, our *in silico* and *in vitro CACNA1A*^+/−^ model have shown two major cellular mechanisms contributing to the *CACNA1A*^+/−^ network phenotype: reduced function and expression of potassium channels leading to increased intrinsic excitability and reduced synaptic function. When attempting to alter one of these mechanisms, by applying the non-selective voltage-gated potassium channel blocker, 4-AP, we only observed a partial rescue of the electrophysiological phenotype. Treatment of episodic ataxia patients with 4-AP alleviates the frequency and severity of episodic attacks, but does not fully prevent them.^39^ This shows a level of predictive validity of our model and the potential of the model for development of therapeutic strategies. One major drawback of this model is that it is based on one genetic background in which we did not mimic a specific human variant. Therefore, we cannot associate our findings with a patient-specific clinical facet of *CACNA1A*-related disorders. We and others have shown that haploinsufficiency of *CACNA1A* is associated with a clinically variable neurodevelopmental phenotype.^5,7^ This indicates that the genetic background may play a role in the clinical presentation of *CACNA1A* haploinsufficiency. Therefore, further investigation, using multiple patient iPSC-derived models is needed.

While excitatory neurons express *CACNA1A* and provide novel mechanistic insights, these neurons might not perfectly reflect all phenotypic features observed in patients, such as cerebellar degeneration. Purkinje cells might reveal additional alterations that are relevant to some of the phenotypic manifestations (ataxia) and response to certain interventions (4-AP). Protocols to differentiate iPSCs towards Purkinje cells are available and have been employed to model spinocerebellar ataxia type 6.^62–67^ Patient iPSC-derived Purkinje cells exhibited vulnerability towards thyroid hormone depletion,^62^ which may be part of a pathological mechanism in spinocerebellar ataxia type 6. Generation of iPSC-derived Purkinje cells with *CACNA1A* haploinsufficiency may therefore provide additional mechanistic insights, which can be correlated to the existing literature on rodent models. With regard to cortical neurons, modeling *CACNA1A* haploinsufficiency with cortical parvalbumin-positive interneurons may also shed light on the pathophysiology of *CACNA1A*-related disorders in the cerebral cortex. Partial or complete loss of Ca_v_2.1 has been shown to impair the presynaptic release of GABA specifically in parvalbumin-positive interneurons.^17,55^ Co-culturing iPSC-derived glutamatergic and GABAergic neurons yields functionally mature interneurons that exhibit inhibitory control on the neuronal network,^34,68,69^ but these protocols still generate few mature parvalbumin-positive interneurons.^34^ Development of new strategies for the differentiation of iPSCs to mature parvalbumin-positive interneurons will boost research into their function in several neurological disorders.

## Abbreviations

AMPA: α-amino-3-hydroxy-5-methyl-4-isoxazole-propionate
DIV: Days in vitro
iPSC: induced pluripotent stem cell
MEA: micro-electrode array
NMDA: N-methyl-D-aspartate.

