## Supplementary material for "*CACNA1A* haploinsufficiency leads to reduced synaptic function and increased intrinsic excitability"

### Supplementary Materials and methods

#### Neuronal differentiation

iPSCs were differentiated into cortical glutamatergic neurons by doxycycline-inducible *Ngn2* overexpression,[1] according to a previously published protocol.[2] At 80-90% confluency (DIV0), single cells were generated from rtTA/*Ngn2*-positive iPSCs for neuronal differentiation by incubating with TrypLE™ Express (Gibco, # 12604021) at 37°C/5% $CO_2$. The iPSCs were resuspended in Essential 8™ Basal medium (Gibco, #A1517001), supplemented with RevitaCell (Gibco, #A2644501), primocin, and doxycycline (4 µg/mL; Sigma Aldrich, #D9891). The plates were pre-coated with poly-L-ornithine hydrobromide (50 µg/mL; Sigma-Aldrich, # P3655) in borate buffer (50 mM) for 3h at 37°C/5% $CO_2$, followed by overnight incubation with Biolaminin 521 LN (Biolamina, LN521) at 4°C. For RNA and western blots, iPSCs were plated at a density of 300,000 and 250,000 cells per well of a 6-well-plate, respectively. For single cell electrophysiology and immunocytochemistry, cells were plated at a density of 20,000 (single cell electrophysiology and synapse density) or 10,000 (morphological reconstruction) cells per well of a 24-well plate. At DIV1, culture medium was changed to DMEM/F12 medium (Gibco, #11320074), supplemented with MEM non-essential amino acid solution (Sigma-Aldrich, #M7145), N2 (Gibco, #17502048), recombinant human BDNF (10 ng/mL; PromoCell, #C-6621), NT3 (10 ng/mL; PromoCell, #C-66425), doxycycline, and mouse laminin from Engelbreth-Holm-Swarm sarcoma (0.2 µg/mL; Sigma-Aldrich, #L2020). To support neuronal maturation and viability, rat embryonic astrocytes were added to the neuronal culture in a 1:1 ratio/well at DIV2. As CACNA1A is expressed in rat astrocytes (Supplementary Fig. 1G), neurons used to detect CACNA1A were cultured without astrocytes and were kept under the same conditions as astrocyte-supported cultures. After three days, the medium was changed to Neurobasal medium (Gibco) supplemented with B-27 (20 µg/mL; Gibco, #17504001), primocin, GlutaMAX (10 µg/mL; Gibco, #35050038), BDNF, NT3, and doxycycline. Furthermore, cytosine β-D-arabinofuranoside hydrochloride (Ara-C) (2 µM; Sigma-Aldrich, #C6645) was added once at DIV3, to remove any proliferating cell from the culture. From DIV6 to DIV9, half of the medium was refreshed every other day with fresh Neurobasal medium supplemented with B-27, primocin, GlutaMAX, BDNF, NT3 and doxycycline. From DIV10 onwards, every other day half of the medium was refreshed with neurobasal medium without doxycycline, and additional 2.5% fetal bovine serum (FBS; Sigma-Aldrich, #F7524), to support astrocyte viability. Throughout the entire differentiation process, the cultures were incubated at 37°C/5% $CO_2$.

#### Immunocytochemistry

For the characterization of the generated induced pluripotent stem cell (iPSC) lines, cells were fixated with 4% paraformaldehyde/4% sucrose for 15 min and permeabilized with 0.2% Triton X-100 (Sigma Aldrich, #T8787) in PBS for 10 min at room temperature. The cells were blocked in blocking buffer containing 5% normal goat serum (Invitrogen, #10000C) in PBS for 1 hr at room temperature. Primary and secondary antibodies (Supplementary Table 3), diluted in blocking buffer, were incubated overnight at 4°C, and for 1 hr at room temperature, respectively. Cells were mounted with fluorescence mounting medium (DAKO, #S302380). Images were captured by a Zeiss Axio Imager Z1.

#### Western blot

For western blot, cell lysates were made from neuronal cultures without astrocytes at DIV21 in order to determine the CACNA1A level. For cell lysis, the culture medium was aspirated, and each well was washed with 2 mL ice-cold DPBS (Gibco, #14040117). Afterwards, lysis buffer (RIPA buffer [pH 7.5; 150 mM NaCl (Sigma-Aldrich), 50 mM Tris-HCl (Invitrogen), 1% Triton X-100, 1 mM EDTA (Sigma-Aldrich)], supplemented with protease inhibitors [cOmplete Mini; Roche] in a ratio 6:1) was added to each well. Protein concentrations were determined by the Pierce™ BCA protein assay (Thermo Fisher Scientific, #23225). Protein samples were loaded equally (10 µg/sample) and separated on 4-15% Mini-PROTEAN TGX

Stain-Free gels (Bio-Rad, #4568084). The proteins were transferred to PVDF membranes (Bio-Rad, #1704156). The membranes were blocked with 5% non-fat dry milk (Santa Cruz Biotechnology, #sc-2325) in 0.1% PBS-T for 1h at room temperature. Primary antibodies (Supplementary Table 3) were diluted in 5% non-fat dry milk for overnight incubation at 4°C. For visualization, the blots were incubated in horseradish peroxidase-conjugated secondary antibody (HRP) for 1h at room temperature (Supplementary Table 3). After incubation, blots were washed and incubated for 5 minutes at room temperature with ECL reagent (SuperSignal™ West Femto Maximum Sensitivity Substrate, Thermo Fisher Scientific, # 34095) and imaged using ChemiDoc™ Touch Gel Imaging System (Bio-Rad). The protein bands were quantified using ImageJ Software. GAPDH was used as loading control.

### Quantification of mRNA by RT-qPCR

Total RNA was isolated and purified with NucleoSpin RNA Mini Kit (Macherey-Nagel, #740955), according to the manufacturer's instructions. The RNA was reverse-transcribed into cDNA using iScript cDNA Synthesis Kit (Bio-Rad, #1708891). Human-specific primers were designed with Primer3plus (Supplementary Table 3). The qPCR was run on a QuantStudio™ 3 Real-Time PCR System (Applied Biosystems) using GoTaq qPCR Master Mix (Promega, #A6002). The qPCR program was designed as followed: after an initial denaturation step at 95 °C for 2 min, PCR amplifications proceeded for 40 cycles of 95 °C for 30 sec and 60 °C for 30 sec, which was followed by a melting curve stage. All samples were analyzed in triplicates in the same run, placed in adjacent wells. Reverse transcriptase-negative controls and no template-controls were included. The arithmetic mean of the Ct values of the technical triplicates was used for calculations. Relative mRNA expression levels were calculated using the $2^{-\Delta\Delta Ct}$ method[3] with normalization to housekeeping genes (*GUSB* and *HPRT1*) and control expression levels.

### Micro-electrode array recordings and data analysis

Recordings of control neuronal networks with 10 µM 1-Naphthyl acetyl spermine trihydrochloride (Naspm; Tocris, #2766) treatment over development (Supplementary Figure 2C-E) were performed on multiwell-MEAs that consisted of 24 wells (Multi Channel Systems, MCS GmbH, Reutlingen, Germany) based on a previously published protocol.[4] In short, each MEA well was embedded with 12 gold electrodes with a diameter of 30 µm, spaced 300 µm apart from each other. Basal neuronal network activity was recorded for 10 minutes every week starting from DIV14 to DIV63, after a 10-minute acclimatization period, in a recording chamber maintained at 37°C and oxygenated with 95% $O_2$/5% $CO_2$. For all recordings, the sampling rate was set at 10 kHz and signals were filtered with a high-pass 2nd order Butterworth filter with a 100 Hz cut-off and a low-pass 4th order Butterworth filter with a 3500 Hz cut-off. The threshold for spike detection was set at ±4.5 standard deviations. Data analysis was performed using Multiwell-Analyzer software (Multi Channel Systems, MCS GmbH, Reutlingen, Germany), and a custom-made code developed in MATLAB (The Mathworks, Natick, MA, USA). To detect bursts, the Multiwell-Analyzer build-in burst detection algorithm was used. A burst was defined when 5 spikes were in close proximity with a maximum inter spike interval (ISI) of 50 ms to start a burst, a maximum ISI of 50 ms to end a burst, and a minimum inter burst interval (IBI) of 100 ms. Network bursts were defined using the single burst detection as an intermediate step.[5] A network burst was defined when the number of distinct bursting channels were at least 50%, and at some point during the sequence, at least 25% of the channels were bursting at the same time. The network burst frequency was derived by dividing all network burst in the recording by the length of the recording in seconds. Mossink *et al.* provides a detailed description of the calculation of other network parameters.[4]

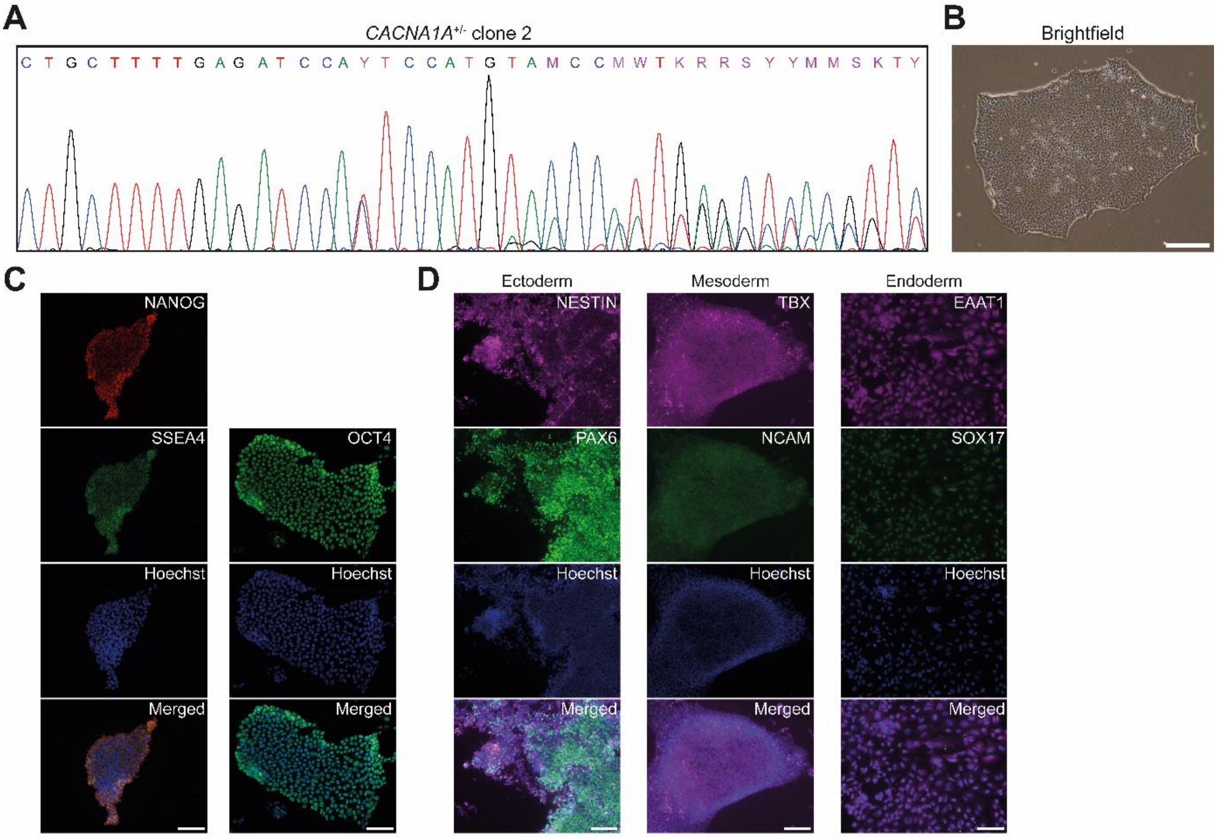


E

| Identified by | Position (GRCh38) | Strand | Sequence | # mismatches | Score | Primers |
|---|---|---|---|---|---|---|
| Benchling | chr22: 32872209 | 1 | ATGTCAGCTCAATGGGAACA | 3 | 1.478 | FW: TTCTCCCACATTGCGTGTAG<br>RV: CTAGGATCGCAGGGAAAGTG |
| Benchling | chr19: 17038479 | 1 | AGGTGAGCTGAGTGGGTACA | 3 | 1.4 | FW: TTGGAGCTAAGCCTGGAGAC<br>RV: CACTTGGGGGCTATTTATGC |
| Benchling | chr9: 5336494 | -1 | AACGGAGCTGAATGGGTACA | 4 | 1.377 | FW: CAGCAGGAGAAGGAAATCTCTG<br>RV: AGAAGAAGAGCAGCAAGACCTG |
| CRISPOR | chr8: 40177980 | 1 | ACGTAAGCTCAGTGGGTACA | 2 | 0.62 | FW: CCCTGAGGAGTGAATCAGATG<br>RV: TCAGCAAATTGCAGGGTATG |
| CRISPOR | chr1: 175678334 | -1 | ATGTGAATTCAGTGGGTACA | 4 | 0.57 | FW: TTTCTTACCCTCAGCCATCC<br>RV: ACCTCTGCCTCTGGACTTTG |
| CRISPOR | chr9: 137955772 | 1 | GCGAGAGCTCAACGGGTACC | 4 | 0.08 | FW: TTTGCCAAGGAGCGAGAGAG<br>RV: ATGCCCCATGTGGAGGAAAG |

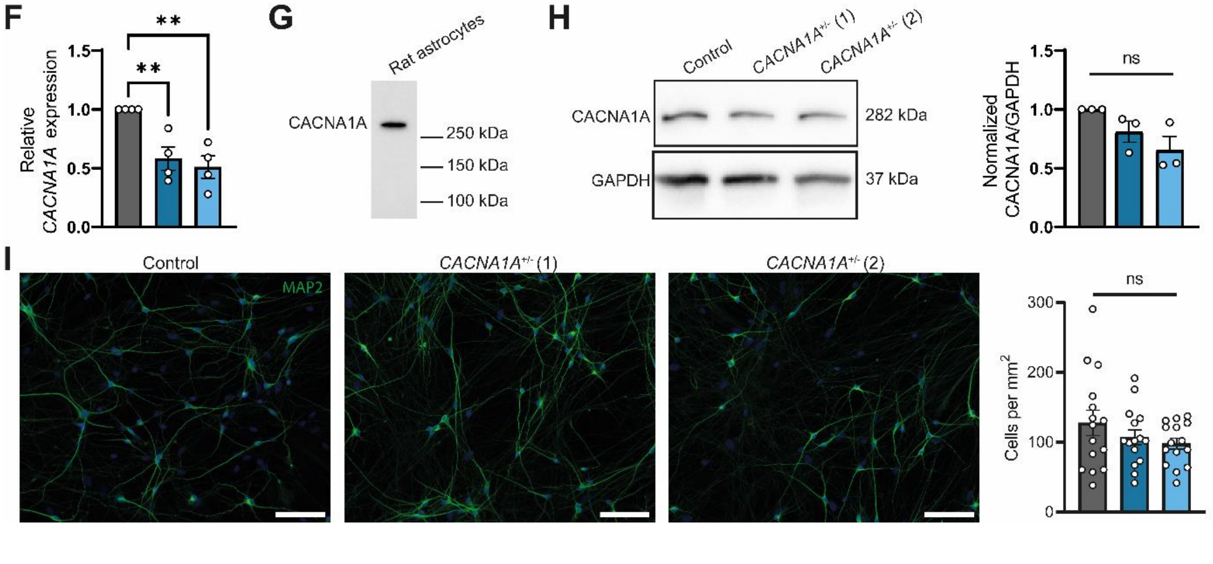

**Supplementary Figure 1 Characterization of *CACNA1A*$^{+/-}$ cell lines.** (**A**) Chromatogram of sequencing result from *CACNA1A*$^{+/-}$ (2) (insertion of 1 nucleotide). (**B**) Brightfield image of a *CACNA1A*$^{+/-}$ (2) colony, showing typical iPSC-like morphology (scale bar 200 μm). (**C**) Representative fluorescent images of *CACNA1A*$^{+/-}$ (2) colonies showing expression of pluripotency markers NANOG, SSEA4 and OCT4 (scale bar 100 μm). (**D**) Representative fluorescent images of *CACNA1A*$^{+/-}$ (2) colonies differentiated to the three germ layers showing expression of ectodermal markers NESTIN and PAX6, mesodermal makers TBX and NCAM, and endodermal markers EAAT1 and SOX17 (scale bar 100 μm). (**E**) Top six potential off-target sites as predicted by Benchling (Biology Software) and CRISPOR[6] have been sequenced and no off-target editing has been detected. (**F**) Normalized *CACNA1A* expression at DIV49. $n = 4$ for all cell lines. (**G**) Western Blot showing CACNA1A protein in rat astrocytes. (**H**) Representative Western Blot and quantification of CACNA1A protein levels normalized to GAPDH and control levels in DIV21 neurons grown without astrocytes. $n = 3$ for all cell lines. (**I**) Representative images of MAP2-positive neurons at DIV21 (scale bar 100 μm), and quantification of MAP2-positive cells per $mm^2$. $n = 15/2$ for all cell lines. $^{**}p < 0.01$, Ordinary one-way ANOVA.

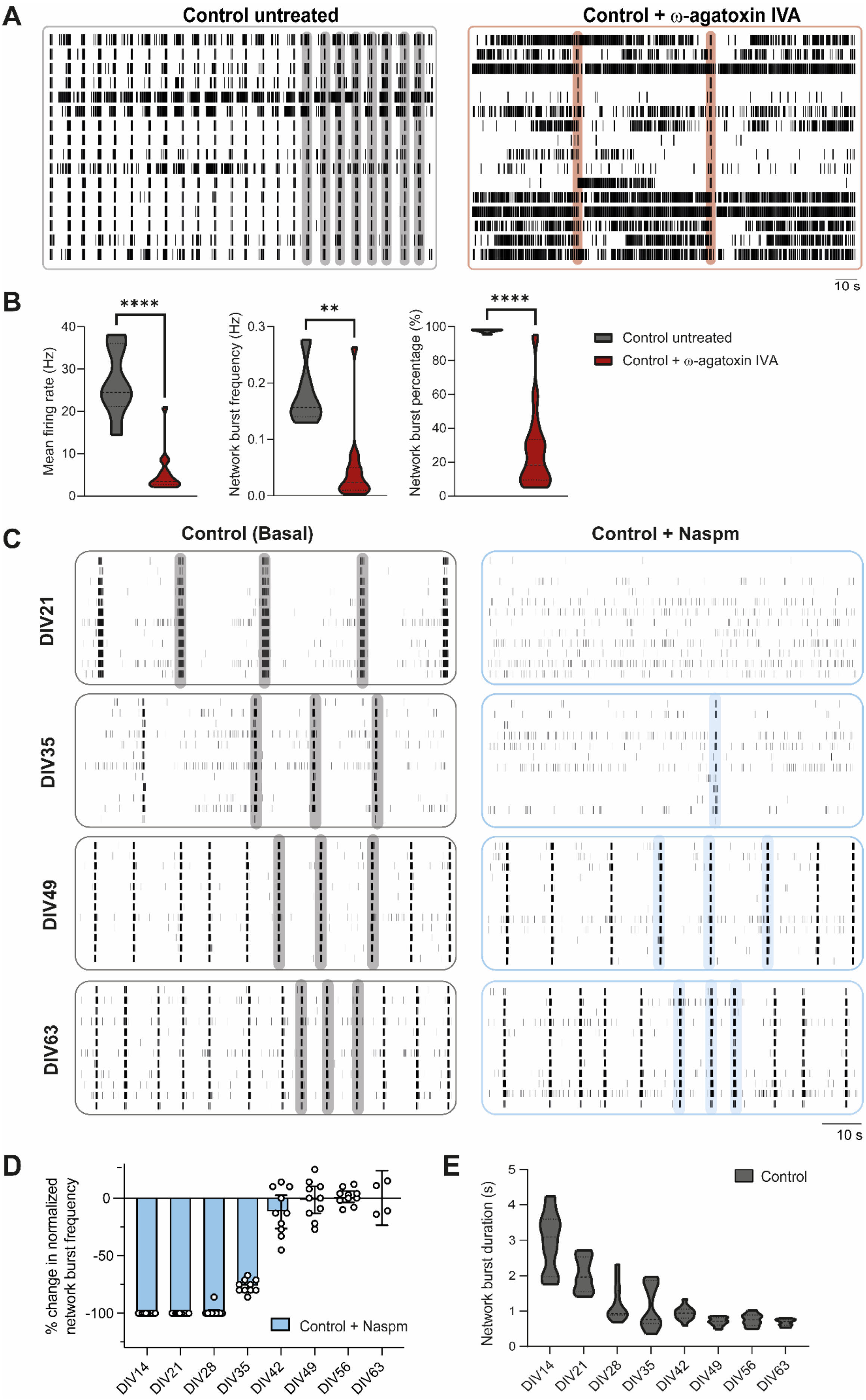

A
Control untreated
Control + ω-agatoxin IVA
10 s
B
****
**
****
Mean firing rate (Hz)
Network burst frequency (Hz)
Network burst percentage (%)
Control untreated
Control + ω-agatoxin IVA
C
Control (Basal)
Control + Naspm
DIV21
DIV35
DIV49
DIV63
10 s
D
% change in normalized network burst frequency
Control + Naspm
DIV14
DIV21
DIV28
DIV35
DIV42
DIV49
DIV56
DIV63
E
Network burst duration (s)
Control

**Supplementary Figure 2 Effects of $Ca_v2.1$ and GluA2-lacking AMPA receptor blockage on control neuronal networks.** (**A**) Representative 3-min rasterplots of spontaneous activity from a control neuronal network at DIV51 either untreated or treated with 200 nM ω-agatoxin IVA. (**B**) Quantification of network parameters including mean firing rate, network burst frequency and network burst percentage (percentage of spikes within a network burst). $n = 6$ for Control untreated; $n = 15$ for Control + ω-agatoxin IVA. Dashed line represents the median, dotted line represents the quartiles. $^{**}p < 0.01$, $^{****}p < 0.0001$, Mann-Whitney test. (**C**) Representative 100-sec rasterplots of spontaneous activity from control neuronal network at DIV21, 35, 49, and 63 as indicated, on basal condition and treated with 10 μM Naspm to block GluA2-lacking AMPA receptors. (**D**) Quantification of change in normalized network burst frequency in the first 100 sec after adding Naspm for control neuronal network recorded throughout development at DIV14-63 as indicated. The values are normalized to the activity 5 min before adding Naspm for each well. $n = 10/2$. Data represent means ± SEM. (**E**) Quantification of network burst duration of control neuronal networks recorded throughout development at DIV14-63 as indicated. $n = 10/2$. Dashed line represents the median, dotted line represents the quartiles.

A

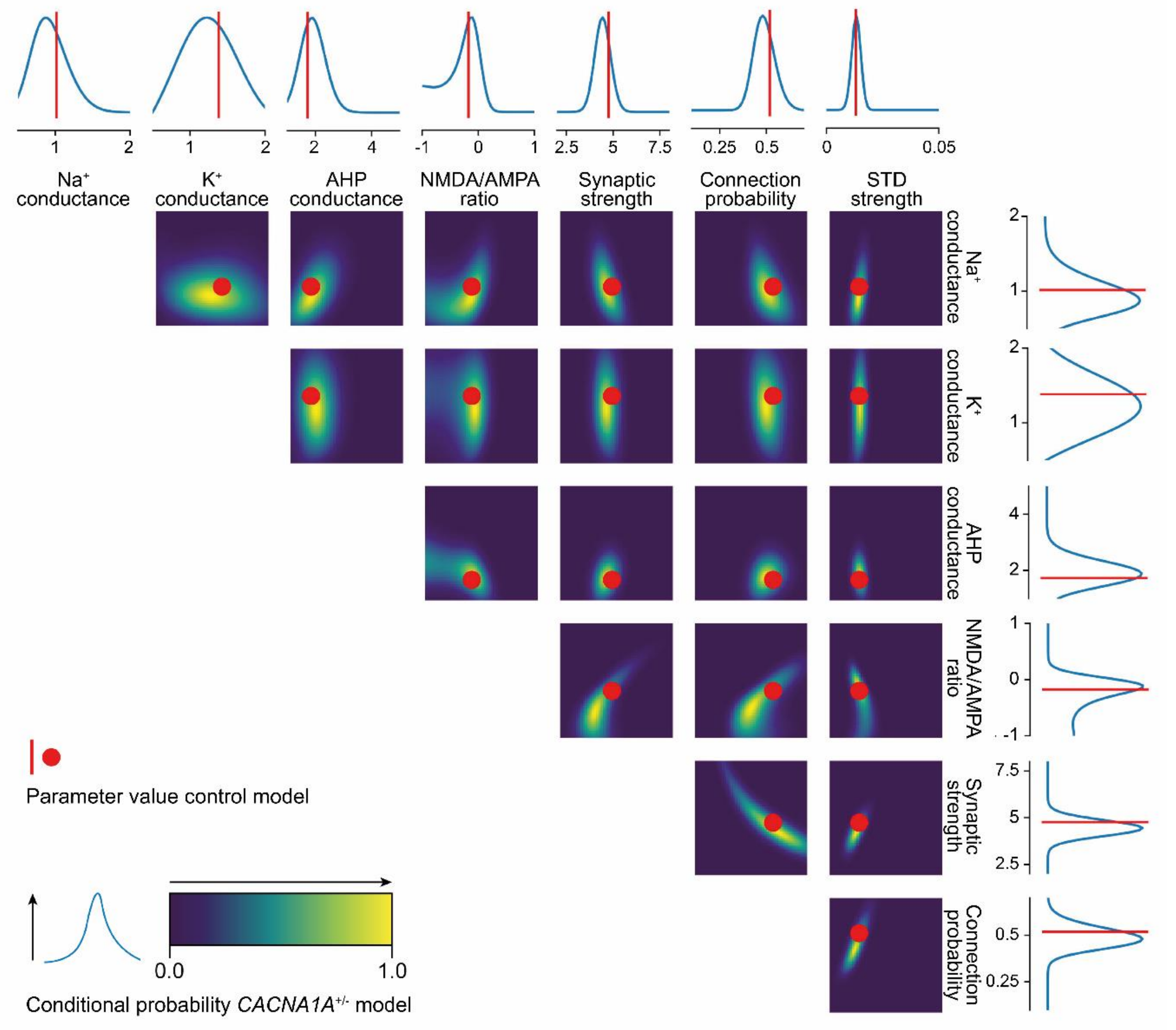
Na+ conductance
K+ conductance
AHP conductance
NMDA/AMPA ratio
Synaptic strength
Connection probability
STD strength
Parameter value control model
0.0
1.0
Conditional probability CACNA1A+/- model

**Supplementary Figure 3 Conditional distribution of *in silico* model parameters.** (**A**) Conditional distribution showing the distribution of parameter values (top and right line plots), or combinations of two parameters (heatmaps), that likely result in *CACNA1A*$^{+/-}$ network activity (blue distribution and clouds), when all other parameters are set to the optimal values to recreate control network activity (red lines and dots).

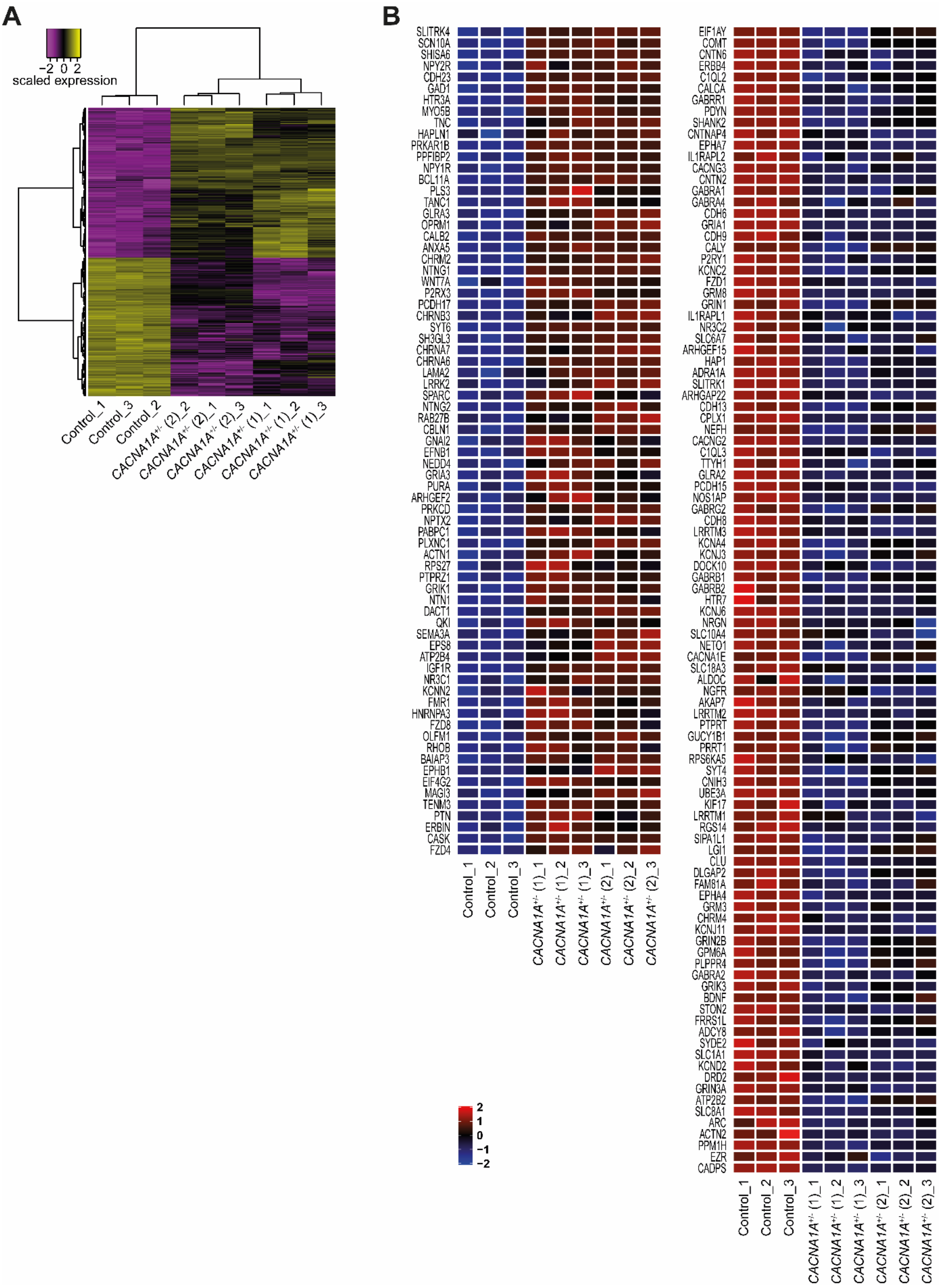
A
−2 0 2
scaled expression
Control_1
Control_3
Control_2
CACNA1A+/- (2)_2
CACNA1A+/- (2)_1
CACNA1A+/- (2)_3
CACNA1A+/- (1)_1
CACNA1A+/- (1)_2
CACNA1A+/- (1)_3
B
SLITRK4
SCN10A
SHISA6
NPY2R
CDH23
GAD1
HTR3A
MYO5B
TNC
HAPLN1
PRKAR1B
PPFIBP2
NPY1R
BCL11A
PLS3
TANC1
GLRA3
OPRM1
CALB2
ANXA5
CHRM2
NTNG1
WNT7A
P2RX3
PCDH17
CHRNB3
SYT6
SH3GL3
CHRNA7
CHRNA6
LAMA2
LRRK2
SPARC
NTNG2
RAB27B
CBLN1
GNAI2
EFNB1
NEDD4
GRIA3
PURA
ARHGEF2
PRKCD
NPTX2
PABPC1
PLXNC1
ACTN1
RPS27
PTPRZ1
GRIK1
NTN1
DACT1
QKI
SEMA3A
EPS8
ATP2B4
IGF1R
NR3C1
KCNN2
FMR1
HNRNPA3
FZD8
OLFM1
RHOB
BAIAP3
EPHB1
EIF4G2
MAGI3
TENM3
PTN
ERBIN
CASK
FZD4
Control_1
Control_2
Control_3
CACNA1A+/- (1)_1
CACNA1A+/- (1)_2
CACNA1A+/- (1)_3
CACNA1A+/- (2)_1
CACNA1A+/- (2)_2
CACNA1A+/- (2)_3
2
1
0
−1
−2
EIF1AY
COMT
CNTN6
ERBB4
C1QL2
CALCA
GABRR1
PDYN
SHANK2
CNTNAP4
EPHA7
IL1RAPL2
CACNG3
CNTN2
GABRA1
GABRA4
CDH6
GRIA1
CDH9
CALY
P2RY1
KCNC2
FZD1
GRM8
GRIN1
IL1RAPL1
NR3C2
SLC6A7
ARHGEF15
HAP1
ADRA1A
SLITRK1
ARHGAP22
CDH13
CPLX1
NEFH
CACNG2
C1QL3
TTYH1
GLRA2
PCDH15
NOS1AP
GABRG2
CDH8
LRRTM3
KCNA4
KCNJ3
DOCK10
GABRB1
GABRB2
HTR7
KCNJ6
NRGN
SLC10A4
NETO1
CACNA1E
SLC18A3
ALDOC
NGFR
AKAP7
LRRTM2
PTPRT
GUCY1B1
PRRT1
RPS6KA5
SYT4
CNIH3
UBE3A
KIF17
LRRTM1
RGS14
SIPA1L1
LGI1
CLU
DLGAP2
FAM81A
EPHA4
GRM3
CHRM4
KCNJ11
GRIN2B
GPM6A
PLPPR4
GABRA2
GRIK3
BDNF
STON2
FRRS1L
ADCY8
SYDE2
SLC1A1
KCND2
DRD2
GRIN3A
ATP2B2
SLC8A1
ARC
ACTN2
PPM1H
EZR
CADPS
Control_1
Control_2
Control_3
CACNA1A+/- (1)_1
CACNA1A+/- (1)_2
CACNA1A+/- (1)_3
CACNA1A+/- (2)_1
CACNA1A+/- (2)_2
CACNA1A+/- (2)_3

**Supplementary Figure 4 Transcriptional changes in *CACNA1A*$^{+/-}$ neurons.** (**A**) Heatmap depicting scaled expression of differentially expressed genes (DEGs) in the nine samples at DIV49. (**B**) Heatmap showing relative expression (Z scores) of a selection of DEGs that are known to play roles in ion channels and synapses, as exported from SynGO release “20231201”. The scale of the Z scores is indicated from -2 (blue) to 2 (red).

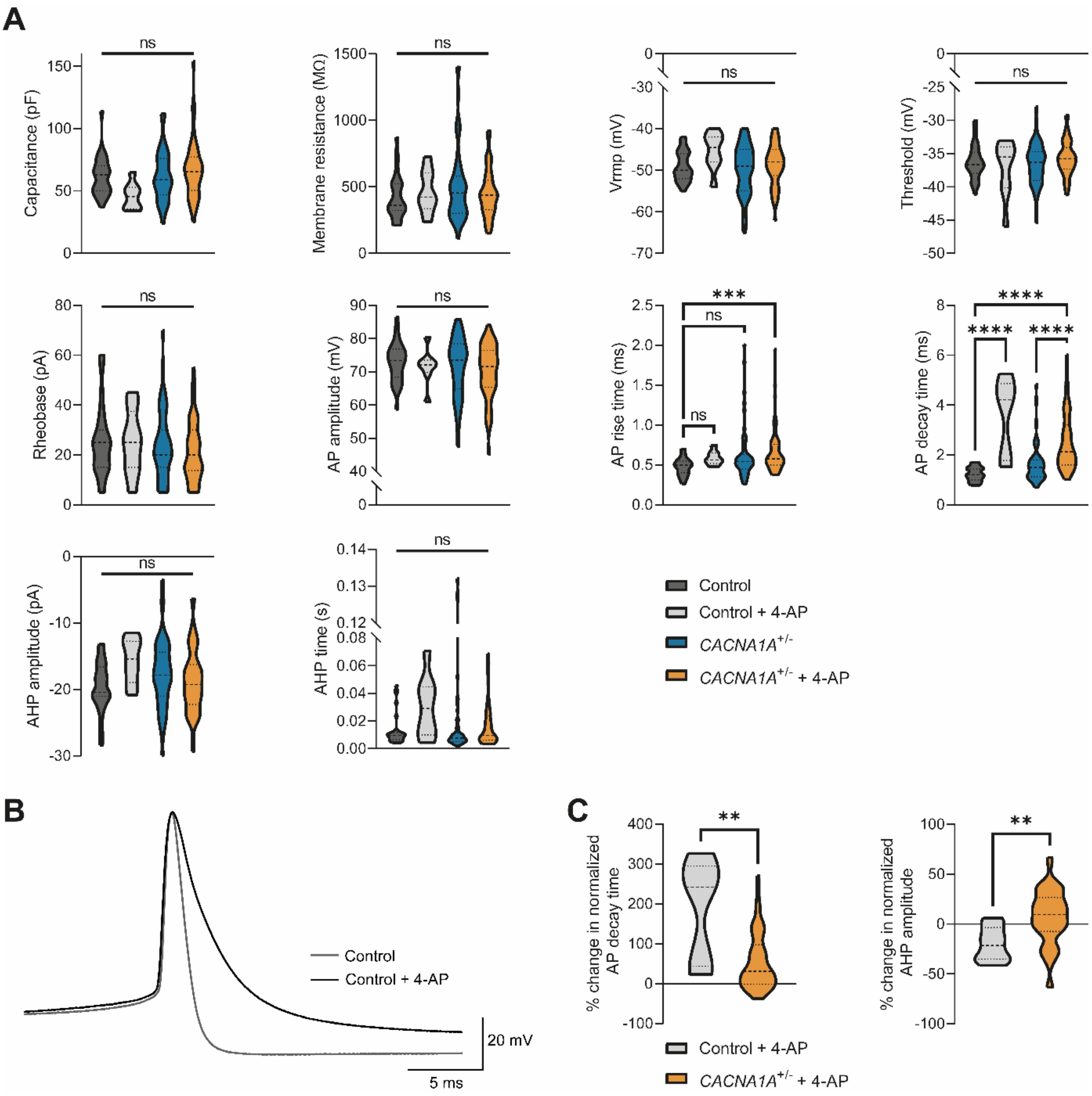

A
Capacitance (pF)
Membrane resistance (MΩ)
Vrmp (mV)
Threshold (mV)
Rheobase (pA)
AP amplitude (mV)
AP rise time (ms)
AP decay time (ms)
AHP amplitude (pA)
AHP time (s)
ns
***
****
Control
Control + 4-AP
CACNA1A+/-
CACNA1A+/- + 4-AP
B
Control
Control + 4-AP
20 mV
5 ms
C
% change in normalized AP decay time
% change in normalized AHP amplitude
**
Control + 4-AP
CACNA1A+/- + 4-AP

**Supplementary Figure 5 Effect of 4-AP on control and *CACNA1A*$^{+/-}$ neurons.** (**A**) Quantification of passive and active intrinsic properties as indicated. *n = 31/3* for Control; *n = 8/2* for Control + 4-AP; *n = 55/3* for *CACNA1A*$^{+/-}$ ((1): *25/3* and (2): *30/3*); *n = 58* for *CACNA1A*$^{+/-}$ + 4-AP ((1): *30/3* and (2): *28/3*). Dashed line represents the median, dotted line represents the quartiles. *$p < 0.05$, ***$p < 0.001$, ****$p < 0.0001$, Kruskal-Wallis test with post hoc Dunn's correction. (**B**) Representative outline of single action potential waveforms of control and 4-AP-treated control neurons at DIV42. (**C**) Quantification of change in normalized decay time and afterhyperpolarization (AHP) amplitude in 4-AP-treated control and *CACNA1A*$^{+/-}$ neurons at DIV42. Values of the decay time and AHP amplitude were normalized to the means of the untreated condition. *n = 8/2* for Control + 4-AP; *n = 58* for *CACNA1A*$^{+/-}$ + 4-AP ((1): *30/3* and (2): *28/3*). Dashed line represents the median, dotted line represents the quartiles. **$p < 0.01$, Mann-Whitney test. Abbreviations: AP = action potential; Vrmp = resting membrane potential.

**Supplementary Table 1 Description of *in silico* model parameters**

| Model parameter | Description |
|---|---|
| Sodium conductance | Maximum conductance of the neuronal membrane to sodium ions as a result of voltage-gated sodium channels. Scales with the amount and the maximum opening probability of these channels. |
| Potassium conductance | Maximum conductance of the neuronal membrane to potassium ions as a result of delayed rectifier potassium channels. Scales with the amount and the maximum opening probability of these channels. |
| sAHP conductance | Maximum conductance of the neuronal membrane to potassium ions as a result of slow after-hyperpolarizing potassium channels. Scales with the amount and the maximum opening probability of these channels. The opening probability of these channels increases with every elicited action potential. |
| NMDA/AMPA ratio | The ratio between the amount of NMDA and AMPA postsynaptic current. Changes with the amount and strength of NMDA and AMPA channels. |
| Synaptic Strength | Scales the maximum amplitude of all post-synaptic currents. |
| Connection probability | The probability that two neurons connect via a synapse. Scales with the number of synapses. |
| STD strength | The strength of short-term synaptic depression (STD). A higher value means a more severe depletion of neurotransmitters upon arrival of a pre-synaptic action potential. |

Abbreviations: sAHP = slow afterhyperpolarization; STD = short term depression.

**Supplementary Table 2 Structure matrix based on discriminant analysis of control, *CACNA1A*$^{+/-}$, and 4-AP-treated *CACNA1A*$^{+/-}$ networks**

| Parameter | Function 1 | Function 2 |
|---|---|---|
| Average Network Burst Fano Factor | 0.175* | 0.093 |
| Number of Spikes per Burst | 0.152* | 0.008 |
| Number of Spikes per Network Burst | 0.126* | 0.023 |
| Mean Firing rate | 0.124* | -0.039 |
| ISI CoV | 0.100* | 0.015 |
| Mean ISI within Burst | -0.096* | -0.083 |
| Full Width at Half Height of Normalized Cross Correlation | -0.086* | -0.083 |
| Time to Burst Peak | -0.177 | 0.271* |
| Average Network Burst CoV2 | 0.119 | 0.162* |
| Area Under Normalized Cross Correlation | 0.124 | 0.125* |
| ISI CoV within Network Burst | 0.075 | 0.115* |
| Network Burst Frequency | 0.031 | -0.106* |
| Inter Burst Interval CoV | -0.043 | -0.105* |
| Network Burst Percentage | 0.071 | 0.099* |
| Inter Burst Interval | -0.023 | 0.097* |
| Network Burst Duration | -0.005 | -0.065* |
| Network ISI CoV | 0.027 | -0.062* |
| Burst Frequency | 0.009 | -0.059* |

* Largest absolute correlation between each variable and any discriminant function
Abbreviations: CoV = Coefficient of variation; ISI = Inter spike interval.

**Supplementary Table 3 Antibodies and qPCR primers used in this study**

| **Antibody** | **Dilution** | **Company, Catalog number** |
|---|---|---|
| Rabbit anti-CACNA1A | 1:5000 | Abcam, ab181371 |
| Rabbit anti-GAPDH | 1:2000 | Cell Signaling Technology, 2118L |
| Rabbit anti-NANOG | 1:1000 | Abcam, ab21624 |
| Mouse anti-SSEA4 | 1:100 | Abcam, ab16287 |
| Rabbit anti-OCT4 | 1:200 | Abcam, ab19857 |
| Mouse anti-NESTIN | 1:100 | Thermo Fisher Scientific, MA1-110 |
| Rabbit anti-PAX6 | 1:300 | BioLegend, 901301 |
| Rabbit anti-TBX | 1:200 | LifeSpan Bio, LS-C416020 |
| Mouse anti-NCAM | 1:100 | LifeSpan Bio, LS-C171208 |
| Rabbit anti-EAAT1 | 1:100 | Abcam, ab416 |
| Mouse anti-SOX17 | 1:200 | Genetex, GTX83580 |
| Goat anti-rabbit HRP | 1:50,000 | Invitrogen, G-21234 |
| Goat anti-rabbit Alexa Fluor 647 | 1:1000 | Invitrogen, A-21245 |
| Goat anti-mouse Alexa Fluor 488 | 1:1000 | Invitrogen, A-11029 |
| Goat anti-mouse Alexa Fluor 568 | 1:1000 | Invitrogen, A-11031 |
| Goat anti-rabbit Alexa Fluor 488 | 1:500 | Invitrogen, A-11034 |
| Goat anti-mouse Alexa Fluro 647 | 1:1000 | Invitrogen, A-21236 |
| **qPCR primer target** | **Forward (5'-3')** | **Revers (5'-3')** |
| *CACNA1A* | CAAACTTCTCCGTCAGGGTTAC | CACGTCGATGCCAATGTTAC |
| *GUSB* | CTCATTTGGAATTTTGCCGATT | CCGAGTGAAGATCCCCTTTTTA |
| *HPRT1* | TGACAGTGGCAAAACAATG | GGTCCTTTTCACCAGCAAGCT |

**Supplementary Table 4 Statistical data per figure panel**

| Figure | Parameter per test | I (n) | II (n) | Adjusted p-value |
|---|---|---|---|---|
| Figure 1E | Mann-Whitney | Median Control | Median *CACNA1A*$^{+/-}$ | |
| | ISI CoV within network burst | 3.232 (24) | 2.428 (57) | <0.0001 (****) |
| | Time to burst peak | 303.5 (24) | 407.5 (57) | <0.0001 (****) |
| | Mean firing rate | 15.07 (24) | 6.526 (57) | <0.9999 (ns) |
| | Network burst frequency | 0.1017 (24) | 0.07667 (57) | 0.0396 (*) |
| | Network burst duration | 1.074 (24) | 1.183 (57) | 0.0072 (**) |
| | Network burst percentage | 96.00 (24) | 91.10 (57) | 0.0036 (**) |
| Figure 2B | Two-way ANOVA with Bonferroni correction | Predicted mean *in vitro* | Predicted mean *in silico* | |
| | Mean firing rate | 23.40 (58) | 25.92 (58) | 0.1435 (ns) |
| | Network burst frequency | 0.08795 (58) | 0.10073 (58) | 0.0808 (ns) |
| | Network burst duration | 0.09409 (58) | 0.9259 (58) | 0.9198 (ns) |
| | Network burst percentage | 88.74 (58) | 89.95 (58) | 0.6175 (ns) |
| Figure 3B | Mann-Whitney | Median Control | Median *CACNA1A*$^{+/-}$ | |
| | Synapsin1$^{+}$ punctae per 10 μm | 3.531 (20) | 1.253 (20) | <0.0001 (****) |
| | Homer1$^{+}$ punctae per 10 μm | 3.354 (20) | 1.154 (20) | <0.0001 (****) |
| | Synapsin1$^{+}$/Homer1$^{+}$ punctae per 10 μm | 2.200 (20) | 0.7776 (20) | <0.0001 (****) |
| Figure 3D | Mann-Whitney | Median Control | Median *CACNA1A*$^{+/-}$ | |
| | Number of primary dendrites | 5 (39) | 6 (92) | 0.1348 (ns) |
| | Soma area | 194.8 (39) | 265.4 (92) | 0.0032 (**) |
| | Number of dendritic ends | 11 (39) | 11 (92) | >0.9999 (ns) |
| | Total dendritic length | 850.8 (39) | 991.3 (92) | 0.2228 (ns) |
| Figure 3F | Two-way ANOVA with Bonferroni correction | Predicted mean Control | Predicted mean *CACNA1A*$^{+/-}$ | |
| | Total dendritic length per 10 μm | 18.79 (39) | 22.50 (91) | <0.0001 (****) |
| Figure 3H | Mann-Whitney | Median Control | Median *CACNA1A*$^{+/-}$ | |
| | Burst rate | 2.7 (37) | 0.4 (57) | <0.0001 (****) |
| | Burst duration | 0.3116 (36) | 0.4489 (35) | 0.0306 (*) |
| | Burst max amplitude | 1360 (36) | 626.5 (40) | <0.0001 (****) |
| Figure 4B | Two-way ANOVA with Bonferroni correction | Predicted mean Control | Predicted mean *CACNA1A*$^{+/-}$ | |
| | Normalized network burst frequency | 0.0802 (10) | 0.6926 (24) | <0.0001 (****) |
| Figure 4C | Mann-Whitney | Median Control | Median *CACNA1A*$^{+/-}$ | |
| | Change in normalized network burst frequency | -18.44 (10) | -61.69 (24) | <0.0001 (****) |

| | | | | |
|---|---|---|---|---|
| Figure 4D | Mann-Whitney | Median Control | Median *CACNA1A*$^{+/-}$ | |
| | Percentage of intrinsic activity | 6.793 (10) | 16.74 (23) | 0.0002 (***) |
| Figure 5D | Mann-Whitney | Median Control | Median *CACNA1A*$^{+/-}$ | |
| | Capacitance | 55 (37) | 48 (63) | >0.9999 (ns) |
| | Membrane resistance | 320 (37) | 405 (63) | 0.1380 (ns) |
| | Vrmp | -52 (36) | -50 (60) | >0.9999 (ns) |
| | Threshold | -36.85 (37) | -36.74 (63) | >0.9999 (ns) |
| | Rheobase | 30 (37) | 20 (63) | 0.0050 (**) |
| | AP Amplitude | 66.46 (37) | 67.62 (63) | >0.9999 (ns) |
| | AP Rise time | 0.4382 (37) | 0.4961(63) | 0.4450 (ns) |
| | AP Decay time | 0.9768 (37) | 1.149 (63) | 0.0120 (*) |
| | AHP amplitude | -19.83 (37) | -17.27 (62) | 0.0430 (*) |
| | AHP time | 0.0072 (37) | 0.0070 (61) | >0.9999 (ns) |
| Figure 5E | Two-way ANOVA with Bonferroni correction | Predicted mean Control | Predicted mean *CACNA1A*$^{+/-}$ | |
| | Number of action potentials | 8.838 (37) | 12.91 (63) | <0.0001 (****) |
| Figure 7C | One-way ANOVA with Bonferroni correction | Mean I | Mean II | |
| | ISI CoV within network burst | | | 0.0118 (*) |
| | Control - *CACNA1A*$^{+/-}$ | 3.897 (3) | 2.766 (6) | 0.0494 (*) |
| | Control - *CACNA1A*$^{+/-}$ + 4-AP | 3.897 (3) | 2.539 (22) | 0.0045 (**) |
| | *CACNA1A*$^{+/-}$ - *CACNA1A*$^{+/-}$ + 4-AP | 2.766 (6) | 2.539 (22) | >0.9999 (ns) |
| | Time to burst peak | | | <0.0001 (****) |
| | Control - *CACNA1A*$^{+/-}$ | 214.5 (3) | 315.2 (22) | <0.0001 (****) |
| | Control - *CACNA1A*$^{+/-}$ + 4-AP | 214.5 (3) | 233.9 (22) | 0.5316 (ns) |
| | *CACNA1A*$^{+/-}$ - *CACNA1A*$^{+/-}$ + 4-AP | 315.2 (6) | 233.9 (22) | <0.0001 (****) |
| Figure 7E | Kruskal-Wallis with Dunn's correction | Median I | Median II | |
| | AP Rise time | | | 0.0009 (***) |
| | Control - *CACNA1A*$^{+/-}$ | 0.4969 (31) | 0.5417 (55) | 0.0473 (*) |
| | Control - *CACNA1A*$^{+/-}$ + 4-AP | 0.4969 (31) | 0.5754 (58) | 0.0002 (***) |
| | *CACNA1A*$^{+/-}$ - *CACNA1A*$^{+/-}$ + 4-AP | 0.5417 (55) | 0.5754 (58) | 0.2018 (ns) |
| | AP Decay time | | | <0.0001 (****) |
| | Control - *CACNA1A*$^{+/-}$ | 1.211 (31) | 1.497 (55) | 0.0378 (*) |
| | Control - *CACNA1A*$^{+/-}$ + 4-AP | 1.211 (31) | 2.121 (58) | <0.0001 (****) |
| | *CACNA1A*$^{+/-}$ - *CACNA1A*$^{+/-}$ + 4-AP | 1.497 (55) | 2.121 (58) | <0.0001 (****) |
| | AHP amplitude | | | 0.3945 (ns) |
| | Control | -20.45 (31) | | |

| | | | | |
|---|---|---|---|---|
| | *CACNA1A*$^{+/-}$ | -17.84 (55) | | |
| | *CACNA1A*$^{+/-}$ + 4-AP | -19.27 (58) | | |
| Supplementary Figure 1F | One-way ANOVA with Bonferroni correction | Mean I | Mean II | |
| | Relative *CACNA1A* expression | | | 0.0036 (**) |
| | Control - *CACNA1A*$^{+/-}$ (1) | 1 (4) | 0.5814 (4) | 0.0094 (**) |
| | Control - *CACNA1A*$^{+/-}$ (2) | 1 (4) | 0.5089 (4) | 0.0036 (**) |
| Supplementary Figure 1H | One-way ANOVA with Bonferroni correction | Mean I | Mean II | |
| | Normalized CACNA1A/GAPDH | | | 0.0750 (ns) |
| | Control - *CACNA1A*$^{+/-}$ (1) | 1 (3) | 0.8099 (3) | |
| | Control - *CACNA1A*$^{+/-}$ (2) | 1 (3) | 0.6526 (3) | |
| Supplementary Figure 1I | One-way ANOVA with Bonferroni correction | Mean I | Mean II | |
| | Cells per $mm^2$ | | | 0.2648 (ns) |
| | Control - *CACNA1A*$^{+/-}$ (1) | 127.7 (15) | 107.1 (15) | |
| | Control - *CACNA1A*$^{+/-}$ (2) | 127.7 (15) | 97.94 (15) | |
| Supplementary Figure 2B | Mann-Whitney | Median Control | Median Control + ω-agatoxin IVA | |
| | Mean firing rate | 24.48 (6) | 3.445 (15) | <0.0001 (****) |
| | Network burst frequency | 0.1567 (6) | 0.02333 (15) | 0.0021 (**) |
| | Network burst percentage | 97.83 (6) | 18.26 (15) | <0.0001 (****) |
| Supplementary Figure 4A | Kruskal-Wallis with Dunn's correction | Median I | Median II | |
| | Capacitance | | | 0.2850 (ns) |
| | Control | 63 (31) | | |
| | Control + 4-AP | 45.5 (8) | | |
| | *CACNA1A*$^{+/-}$ | 59 (55) | | |
| | *CACNA1A*$^{+/-}$ + 4-AP | 65.5 (58) | | |
| | Membrane resistance | | | >0.9999 (ns) |
| | Control | 360.0 (31) | | |
| | Control + 4-AP | 423.5 (8) | | |
| | *CACNA1A*$^{+/-}$ | 453.0 (55) | | |
| | *CACNA1A*$^{+/-}$ + 4-AP | 437.0 (58) | | |
| | Vrmp | | | >0.9999 (ns) |
| | Control | -50.0 (31) | | |
| | Control + 4-AP | -44.5 (8) | | |

| | | | |
|---|---|---|---|
| *CACNA1A*$^{+/-}$ | -49.0 (55) | | |
| *CACNA1A*$^{+/-}$ + 4-AP | -48.0 (58) | | |
| Threshold | | | >0.9999 (ns) |
| Control | -36.66 (31) | | |
| Control + 4-AP | -35.49 (8) | | |
| *CACNA1A*$^{+/-}$ | -36.30 (55) | | |
| *CACNA1A*$^{+/-}$ + 4-AP | -35.75 (58) | | |
| Rheobase | | | >0.9999 (ns) |
| Control | 25 (31) | | |
| Control + 4-AP | 25 (8) | | |
| *CACNA1A*$^{+/-}$ | 20 (55) | | |
| *CACNA1A*$^{+/-}$ + 4-AP | 20 (58) | | |
| AP Amplitude | | | >0.9999 (ns) |
| Control | 73.38 (31) | | |
| Control + 4-AP | 72.07 (8) | | |
| *CACNA1A*$^{+/-}$ | 73.52 (55) | | |
| *CACNA1A*$^{+/-}$ + 4-AP | 71.56 (58) | | |
| AP Rise time | | | 0.0070 (**) |
| Control - Control + 4-AP | 0.4969 (31) | 0.5616 (8) | 0.2068 (ns) |
| Control - *CACNA1A*$^{+/-}$ | 0.4969 (31) | 0.5417 (55) | 0.0867 (ns) |
| Control - *CACNA1A*$^{+/-}$ + 4-AP | 0.4969 (31) | 0.5754 (58) | 0.0003 (***) |
| Control + 4-AP - *CACNA1A*$^{+/-}$ | 0.5616 (8) | 0.5417 (55) | >0.9999 (ns) |
| Control + 4-AP - *CACNA1A*$^{+/-}$ + 4-AP | 0.5616 (8) | 0.5754 (58) | >0.9999 (ns) |
| *CACNA1A*$^{+/-}$ - *CACNA1A*$^{+/-}$ + 4-AP | 0.5417 (55) | 0.5754 (58) | 0.3800 (ns) |
| AP Decay time | | | <0.0001 (****) |
| Control - Control + 4-AP | 1.211 (31) | 4.214 (8) | <0.0001 (****) |
| Control - *CACNA1A*$^{+/-}$ | 1.211 (31) | 1.497 (55) | 0.0857 (ns) |
| Control - *CACNA1A*$^{+/-}$ + 4-AP | 1.211 (31) | 2.121 (58) | <0.0001 (****) |
| Control + 4-AP - *CACNA1A*$^{+/-}$ | 4.214 (8) | 1.497 (55) | 0.0016 (**) |
| Control + 4-AP - *CACNA1A*$^{+/-}$ + 4-AP | 4.214 (8) | 2.121 (58) | >0.9999 (ns) |
| *CACNA1A*$^{+/-}$ - *CACNA1A*$^{+/-}$ + 4-AP | 1.497 (55) | 2.121 (58) | <0.0001 (****) |
| AHP amplitude | | | 0.4740 (ns) |
| Control | -20.45 (31) | | |
| Control + 4-AP | -15.42 (8) | | |
| *CACNA1A*$^{+/-}$ | -17.84 (55) | | |
| *CACNA1A*$^{+/-}$ + 4-AP | -19.27 (58) | | |
| AHP time | | | 0.3960 (ns) |

| | | | | |
|---|---|---|---|---|
| | Control | 0.0095 (31) | | |
| | Control + 4-AP | 0.02895 (8) | | |
| | *CACNA1A*$^{+/-}$ | 0.0076 (55) | | |
| | *CACNA1A*$^{+/-}$ + 4-AP | 0.0095 (58) | | |
| Supplementary Figure 4C | Mann-Whitney | Median Control + 4-AP | Median *CACNA1A*$^{+/-}$ + 4-AP | |
| | Change in normalized AP decay time | 242.0 (8) | 30.87 (58) | 0.0042 (**) |
| | Change in normalized AHP amplitude | -21.51 (8) | 9.587 (58) | 0.0068 (**) |

**Supplementary Table 5 Summary data per *CACNA1A*$^{+/-}$ line per figure panel**

| **Figure** | **Parameter** | ***CACNA1A*$^{+/-}$ (1)** | ***CACNA1A*$^{+/-}$ (2)** |
|---|---|---|---|
| Figure 1E | | Median ± IQR (n) | Median ± IQR (n) |
| | ISI CoV within network burst | 2.557 ± 0.579 (29) | 2.393 ± 0.808 (28) |
| | Time to burst peak | 402.5 ± 54 (29) | 411.5 ± 52 (28) |
| | Mean firing rate | 6.166 ± 2.856 (29) | 6.864 ± 8.076 (28) |
| | Network burst frequency | 0.07 ± 0.037 (29) | 0.088 ± 0.053 (28) |
| | Network burst duration | 1.183 ± 0.268 (29) | 1.178 ± 0.321 (28) |
| | Network burst percentage | 91.103 ± 6.5 (29) | 91.93 ± 4.816 (28) |
| Figure 3B | | Median ± IQR (n) | Median ± IQR (n) |
| | Synapsin1$^{+}$ punctae per 10 µm | 2.18 ± 1.62 (10) | 0.95 ± 0.53 (10) |
| | Homer1$^{+}$ punctae per 10 µm | 2.34 ± 1.44 (10) | 0.20 ± 0.97 (10) |
| | Synapsin1$^{+}$/Homer1$^{+}$ punctae per 10 µm | 1.43 ± 1.07 (10) | 0.10 ± 0.79 (10) |
| Figure 3D | | Median ± IQR (n) | Median ± IQR (n) |
| | Number of primary dendrites | 6 ± 3 (45) | 6 ± 2 (47) |
| | Soma area | 250.4 ± 98.8 (45) | 301.5 ± 173.8 (47) |
| | Number of dendritic ends | 12 ± 5 (45) | 10 ± 4 (47) |
| | Total dendritic length | 944.8 ± 595.8 (45) | 1023.4 ± 394.8 (47) |
| Figure 3F | | Predicted mean ± SEM (n) | Predicted mean ± SEM (n) |
| | Total dendritic length per 10 µm | 20.68 ± 0.76 (44) | 24.20 ± 0.73 (47) |
| Figure 3H | | Median ± IQR (n) | Median ± IQR (n) |
| | Burst rate | 0.25 ± 0.65 (34) | 2.1 ± 4.05 (23) |
| | Burst duration | 0.515 ± 0.613 (17) | 0.437 ± 0.360 (18) |
| | Burst max amplitude | 631.3 ± 370.4 (22) | 524.2 ± 401.7 (18) |
| Figure 4B | | Predicted mean ± SEM (n) | Predicted mean ± SEM (n) |
| | Normalized network burst frequency | 0.644 ± 0.029 (12) | 0.742 ± 0.027 (12) |
| Figure 4C | | Median ± IQR (n) | Median (n) |
| | Change in normalized network burst frequency | -88.87 ± 29.83 (12) | -46.01 ± 30.43 (12) |
| Figure 4D | | Median ± IQR (n) | Median (n) |
| | Percentage of intrinsic activity | 16.55 ± 19.65 (12) | 16. 74 ± 20.97 (12) |
| Figure 5D | | Median ± IQR (n) | Median ± IQR (n) |
| | Capacitance | 47 ± 18 (36) | 63 ± 38 (27) |
| | Membrane resistance | 457 ± 306 (36) | 376 ± 195 (27) |
| | Vrmp | -50 ± 9 (36) | -51 ± 6 (26) |
| | Threshold | -36.792 ± 2.997 (36) | -36.133 ± 5.303 (27) |

| | | | |
|---|---|---|---|
| | Rheobase | 20 ± 11 (36) | 20 ± 20 (27) |
| | AP Amplitude | 66.25 ± 10.20 (36) | 70.08 ± 12.07 (27) |
| | AP Rise time | 0.513 ± 0.200 (36) | 0.463 ± 0.237 (27) |
| | AP Decay time | 1.251 ± 0.410 (36) | 1.002 ± 0.436 (27) |
| | AHP amplitude | -16.765 ± 3.692 (34) | -19. 121 ± 7.213 (27) |
| | AHP time | 0.009 ± 0.008 (34) | 0.007 ± 0.003 (27) |
| Figure 5E | | Predicted mean ± SEM (n) | Predicted mean ± SEM (n) |
| | Number of action potentials | 14.0 ± 0.5 (36) | 10.9 ± 0.6 (27) |
| Figure 7C | | Mean ± SEM (n) | Mean ± SEM (n) |
| | ISI CoV within network burst | | |
| | *CACNA1A*$^{+/-}$ | 3.112 ± 0.568 (3) | 2.419 ± 0.468 (3) |
| | *CACNA1A*$^{+/-}$ + 4-AP | 2.881 ± 0.156 (11) | 2.187 ± 0.087 (11) |
| | Time to burst peak | | |
| | *CACNA1A*$^{+/-}$ | 307.8 ± 16.6 (3) | 322.5 ± 2.0 (3) |
| | *CACNA1A*$^{+/-}$ + 4-AP | 244.5 ± 7.2 (11) | 223.2 ± 5.2 (11) |
| Figure 7E | | Median ± IQR (n) | Median ± IQR (n) |
| | AP Rise time | | |
| | *CACNA1A*$^{+/-}$ | 0.580 ± 0.153 (25) | 0.516 ± 0.154 (30) |
| | *CACNA1A*$^{+/-}$ + 4-AP | 0.578 ± 0.309 (29) | 0.573 ± 0.219 (29) |
| | AP Decay time | | |
| | *CACNA1A*$^{+/-}$ | 1.265 ± 0.425 (25) | 1.573 ± 0.692 (30) |
| | *CACNA1A*$^{+/-}$ + 4-AP | 2.137 ± 1.287 (29) | 2.106 ± 2.028 (29) |
| | AHP amplitude | | |
| | *CACNA1A*$^{+/-}$ | -18.615 ± 6.456 (25) | -16.537 ± 5.045 (30) |
| | *CACNA1A*$^{+/-}$ + 4-AP | -17.834 ± 7.359 (29) | -20.407 ± 6.116 (29) |
| Supplementary Figure 4A | | Median ± IQR (n) | Median ± IQR (n) |
| | Capacitance | | |
| | *CACNA1A*$^{+/-}$ | 63 ± 25 (25) | 58 ± 30 (30) |
| | *CACNA1A*$^{+/-}$ + 4-AP | 64 ± 21 (29) | 66 ± 30 (29) |
| | Membrane resistance | | |
| | *CACNA1A*$^{+/-}$ | 467 ± 218 (25) | 433 ± 277 (30) |
| | *CACNA1A*$^{+/-}$ + 4-AP | 417 ± 270 (29) | 439 ± 185 (29) |
| | Vrmp | | |
| | *CACNA1A*$^{+/-}$ | -48 ± 8 (25) | -50 ± 10 (30) |
| | *CACNA1A*$^{+/-}$ + 4-AP | -49 ± 5 (29) | -48 ± 8 (29) |

| | | | |
|---|---|---|---|
| | Threshold | | |
| | *CACNA1A*$^{+/-}$ | -35.623 ± 3.877 (25) | -36.929 ± 3.686 (30) |
| | *CACNA1A*$^{+/-}$ + 4-AP | -36.548 ± 2.624 (29) | -35.639 ±- 3.783 (29) |
| | Rheobase | | |
| | *CACNA1A*$^{+/-}$ | 20 ± 20 (25) | 20 ± 14 (30) |
| | *CACNA1A*$^{+/-}$ + 4-AP | 20 ± 15 (29) | 20 ± 15 (29) |
| | AP Amplitude | | |
| | *CACNA1A*$^{+/-}$ | 70.342 ± 10.978 (25) | 76.072 ± 13.211 (30) |
| | *CACNA1A*$^{+/-}$ + 4-AP | 71.444 ± 9.791 (29) | 73.821 ± 0.219 (29) |
| | AP Rise time | | |
| | *CACNA1A*$^{+/-}$ | 0.580 ± 0.153 (25) | -0.516 ± 0.154 (30) |
| | *CACNA1A*$^{+/-}$ + 4-AP | 0.578 ± 0.309 (29) | 0.573 ± 0.219 (29) |
| | AP Decay time | | |
| | *CACNA1A*$^{+/-}$ | 1.265 ± 0.425 (25) | 1.573 ± 0.692 (30) |
| | *CACNA1A*$^{+/-}$ + 4-AP | 2.137 ± 1.287 (29) | 2.106 ± 2.028 (29) |
| | AHP amplitude | | |
| | *CACNA1A*$^{+/-}$ | -18.615 ± 6.456 (25) | -16.537 ± 5.045 (30) |
| | *CACNA1A*$^{+/-}$ + 4-AP | -17.834 ± 7.359 (29) | -20.407 ± 6.116 (29) |
| | AHP time | | |
| | *CACNA1A*$^{+/-}$ | 0.007 ± 0.003 (25) | 0.009 ± 0.009 (30) |
| | *CACNA1A*$^{+/-}$ + 4-AP | 0.009 ± 0.01 (29) | 0.015 ± 0.018 (29) |
| Supplementary Figure 4C | | Median ± IQR (n) | Median ± IQR (n) |
| | Change in normalized AP decay time | 31.828 ± 79.388 (29) | 29.916 ± 125.10 (29) |
| | Change in normalized AHP amplitude | 1.447 ± 41.868 (29) | 16.082 ± 24.790 (29) |
